# Structural disconnections explain brain network dysfunction after stroke

**DOI:** 10.1101/562165

**Authors:** Joseph C. Griffis, Nicholas V. Metcalf, Maurizio Corbetta, Gordon L. Shulman

## Abstract

Functional connectivity (FC) studies have identified physiological signatures of stroke that correlate with behavior. Using structural and functional MRI data from 114 stroke patients, 24 matched controls, and the Human Connectome Project, we tested the hypothesis that structural disconnection, not damage to critical regions, underlies FC disruptions. Disconnection severity outperformed damage to putative FC connector nodes for explaining reductions in system modularity, and multivariate models based on disconnection outperformed damage models for explaining FC disruptions within and between systems. Across patients, disconnection and FC patterns exhibited a low-dimensional covariance dominated by a single axis linking interhemispheric disconnections to reductions in FC measures of interhemispheric system integration, ipsilesional system segregation, and system modularity, and that correlated with multiple behavioral deficits. These findings clarify the structural basis of FC disruptions in stroke patients and demonstrate a low-dimensional link between perturbations of the structural connectome, disruptions of the functional connectome, and behavioral deficits.

## Introduction

Focal brain lesions cause dysfunction in distributed brain systems. Functional connectivity (FC), which measures the correlation between spontaneous activity fluctuations in remote brain regions (Biswal et al., 1995), has recently been used to identify several systems-scale abnormalities that predict behavioral impairment after stroke (Baldassarre et al., 2016a; Corbetta et al., 2018): reduced interhemispheric system integration (Bauer et al., 2014; Carter et al., 2010; Golestani et al., 2013; He et al., 2007; Lim et al., 2014; van Meer et al., 2012; New et al., 2015; Park et al., 2011; Siegel et al., 2016; Tang et al., 2016; Wang et al., 2010; Wu et al., 2011), reduced ipsilesional system segregation (Baldassarre et al., 2014; Bauer et al., 2012; Eldaief et al., 2016; Ramsey et al., 2016), reduced system modularity (Gratton et al., 2012; Siegel et al., 2018). An important next step is to characterize how the severity of these FC abnormalities depends on the properties of the focal lesion (Carrera and Tononi, 2014; Corbetta et al., 2018).

Most work on this topic has focused on how the severity and form of FC disruptions depends on the FC network properties of damaged grey matter (GM) nodes (Eldaief et al., 2016; Gratton et al., 2012; Nomura et al., 2010; Ovadia-Caro et al., 2013), and broad disruptions of FC and behavior have been proposed to reflect damage to cortical “connector” nodes that interface with multiple systems (Gratton et al., 2012; Warren et al., 2014). However, strokes, tumors, and trauma commonly cause white matter (WM) damage (Corbetta et al., 2015; Esmaeili et al., 2018; Sharp et al., 2014). In stroke, for example, most lesions cause subcortical and WM damage, and pure cortical damage is uncommon (Corbetta et al., 2015). Accordingly, we consider it unlikely that damage to critical GM nodes is the primary factor underlying lesion-induced FC disruptions.

Given that structural connectivity (SC) both directly and indirectly shapes FC in the healthy brain (Adachi et al., 2011; Goni et al., 2014; Greicius et al., 2009; Van Den Heuvel et al., 2009; Honey and Sporns, 2009), we expected that structural disconnection (SDC) plays a similarly fundamental role in determining the severity of FC disruptions caused by stroke. While this expectation aligns with predictions based on simulation studies (Alstott et al., 2009; Cabral et al., 2012; Saenger et al., 2017), empirical support is scarce (Carter et al., 2012) and largely based on evidence from callosal resections and traumatic brain injuries (Jilka et al., 2014; Johnston et al., 2008; Roland et al., 2017). We therefore aimed to test the hypothesis that a stroke’s distributed impact on the structural connectome, not its focal impact on critical FC network nodes, is what determines its impact on FC.

The relationship between SC and FC is an important topic in systems neuroscience (Mišić and Sporns, 2016; Park and Friston, 2013). However, as the structural connectome cannot be experimentally manipulated in human subjects, empirical data relating perturbations of the structural connectome to changes in the functional connectome are scarce (Jilka et al., 2014; Johnston et al., 2008; Roland et al., 2017). Because focal brain lesions can be conceptualized as naturally occurring perturbations of the structural connectome, our second aim was to empirically characterize the relationship between SDC and FC patterns across patients.

We were interested in the dimensionality of this relationship. Stroke produces a small set of related FC abnormalities, and behavioral deficits appear similarly low-dimensional such that a few principal components account for most of the variance in performance within and across cognitive domains (Corbetta et al., 2015). One explanation for this is that strokes often affect multiple proximal fiber pathways within vascular territories (Corbetta et al., 2018). If this is true, then the relationship between SDC and FC patterns should also be low-dimensional, and the FC patterns identified based on their relationships to SDCs should reflect core FC disruptions that have been identified based on their relationships to behavior.

## Results

### Damage and disconnection capture distinct lesion information

Structural MRI data were acquired from 114 sub-acute (i.e. < 2 weeks post-stroke) stroke patients and 24 demographically matched healthy controls (**Table 1**). For each patient, we defined measures of (1) voxel damage, (2) parcel damage, (3) tract SDC, and (4) parcel SDC to quantify the lesion’s structural impact at different spatial scales (**Fig. 1A**, see also **Fig. S1**).

**Table 1.** Demographic information

| **Group** | **Age (Mean/SD)** | **Sex** | **Handedness** | **Education (Mean/SD)** | **Lesion Side** |
| --- | --- | --- | --- | --- | --- |
| Patients (N=114) | 52.4/10.2 | 58 F, 56 M | 106 R, 8 L | 13.3/2.8 | 54 R, 60 L |
| Controls (N=24) | 54.5/13.5 | 12 F, 12 M | 23 R, 1 L | 13.5/2.1 | N/A |

*SD: Standard Deviation, M: Male, F: Female, R: Right, L: Left*

**Figure 1.**
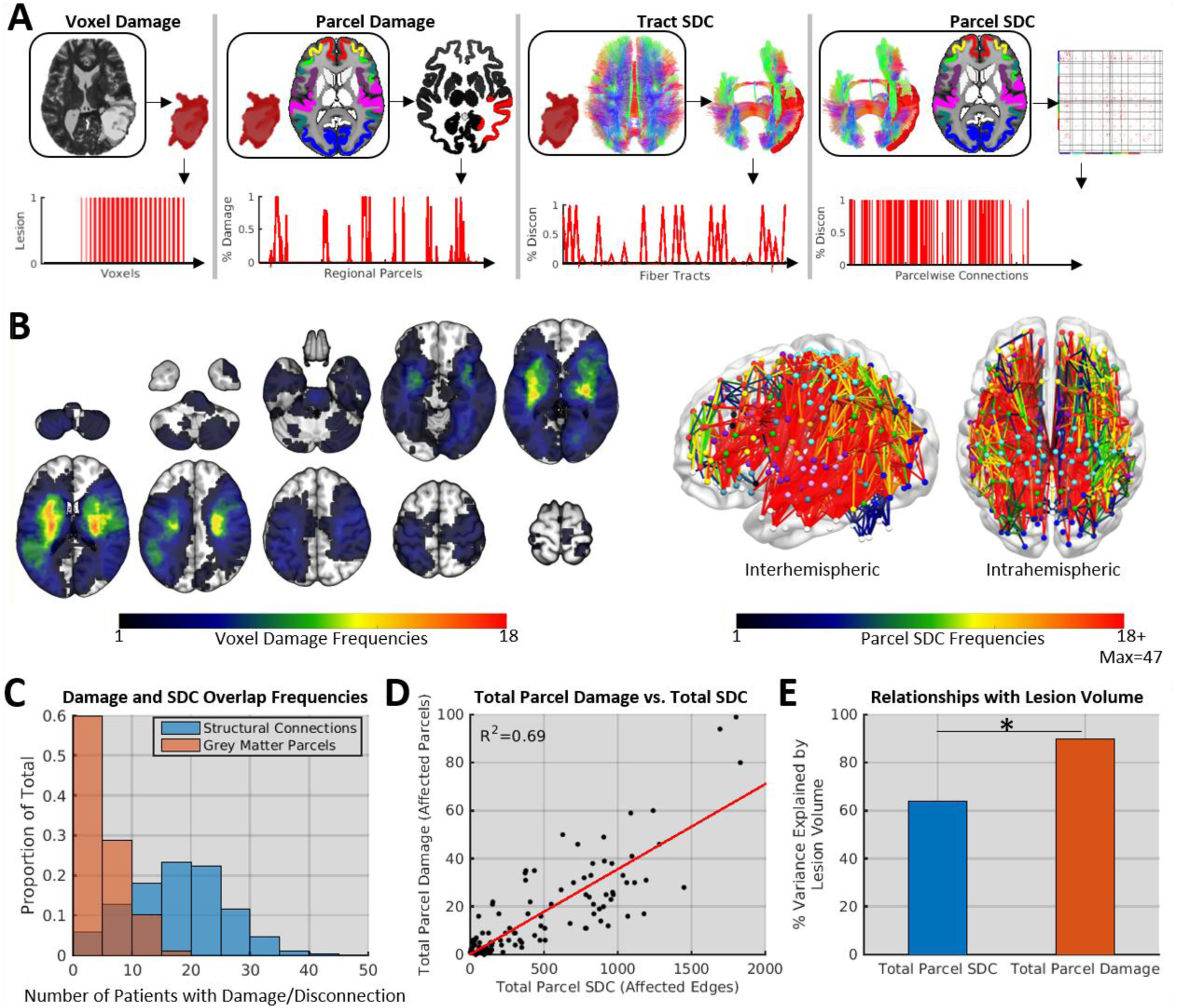
Damage and disconnection data. **A.** Example damage and SDC measures for a single patient. **B.** Group-level voxel damage (left) and parcel SDC (right) topographies. Colors indicate overlap at each voxel/edge. **C.** Group-level parcel SDC and parcel damage overlap frequency distributions. **D.** Relationship between the total number of parcel SDCs and the total number of damaged parcels. **E.** Effects of lesion volume on total parcel SDCs and total parcel damage. *Steiger’s *z*=7.54, *p*<0.001. See also **Figure S1**.

Similar SDCs can be caused by dissimilar lesions due to the spatially distributed nature of SC (Catani et al., 2012). Accordingly, while group-level lesion overlaps were generally infrequent and concentrated in subcortical and WM regions (**Fig. 1B**, left), SDC overlaps were frequent throughout the brain (**Fig. 1B**, right). SDC overlaps were more frequent even when lesion overlap was summarized at the level of whole GM parcels (**Fig. 1C**). The extent of parcel SDCs was only partially explained by the extent of focal damage (**Fig. 1D**). While the number of damaged parcels largely reflected lesion size, lesion size was less strongly related to the number of disconnected parcels (**Fig. 1D**). This is consistent with the idea that a lesion’s impact on the connectome is not entirely dictated by its size, as even small subcortical and WM lesions have the potential to cause widespread SDC (Yeh et al., 2013a).

### Stroke disrupts system-scale functional connectivity

Resting-state fMRI data were used to measure FC between 324 cortical parcels associated with different brain systems (**Fig. 2A-C**). We defined twelve system-scale summary measures to capture reductions of interhemispheric system integration, ipsilesional system segregation, and system modularity. For each patient, we extracted the mean interhemispheric FC values for nine bilateral cortical systems (**Fig. 2D**, left) and averaged across systems to summarize interhemispheric system integration across the cortex (**Fig. 2D**, left inset). We also extracted the mean FC between the ipsilesional DAN and DMN to quantify ipsilesional system segregation (**Fig. 2D**, middle), and we averaged modularity estimates for *a priori* system partitions across multiple edge density thresholds to summarize overall network structure (**Fig. 2D**, right; mean shown in inset).

**Figure 2.**
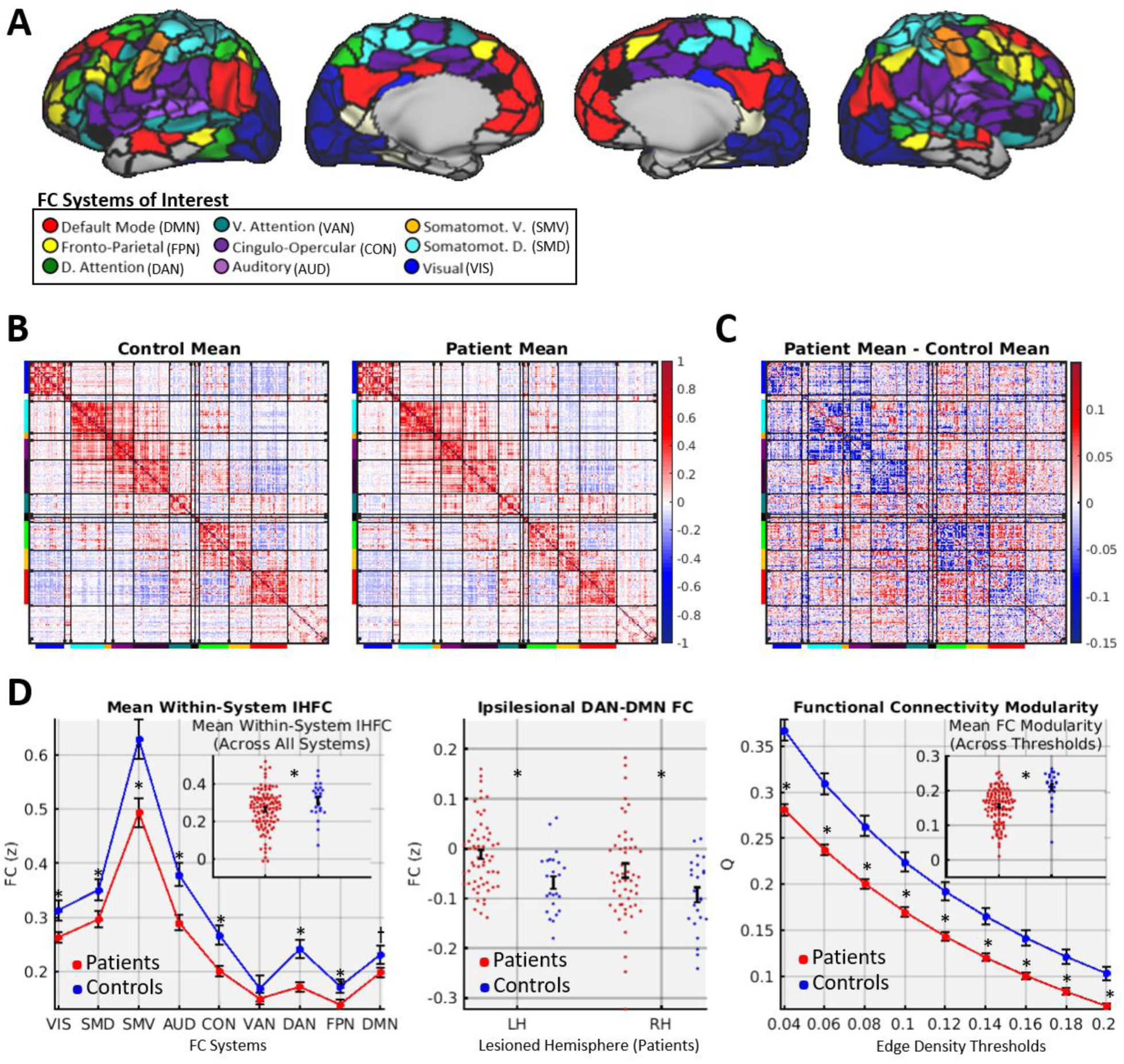
FC disruptions. **A**. Cortical parcels and systems of interest (see also **Fig. S1**). **B.** Mean FC matrices from each group. **C**. Difference in mean matrices. **D.** System-level FC summaries for patients (red) and controls (blue). Plots show means/standard errors. *two-sample *t*-test FDR*p*<0.05.

The mean FC matrices for patients and controls had similar topographies (**Fig. 2B**; *r*=0.96, *p*<0.001), but subtracting the patient matrix from the control matrix revealed magnitude differences that were often in opposite directions for connections with positive vs. negative values in the mean control matrix (**Fig. 2C**; *r*=-0.42, *p*<0.001), consistent with reduced integration within systems and reduced segregation between systems in patients. As expected, patients showed marked abnormalities in FC summary measures of interhemispheric integration, ipsilesional segregation, and system modularity (**Fig. 2D**). These measures were used as dependent variables in subsequent analyses.

### Modularity is better explained by disconnection than damage to critical regions

Damage to parcels with high FC participation coefficients (PC, “connectors”), but not parcels with high FC within-module degree (WMD, “hubs”), has been associated with reduced system modularity (Gratton et al., 2012). However, this effect has not been independently replicated or compared to other plausible mechanisms such as SDC severity. We aimed to replicate this effect in our data and compare its explanatory power to that of a simple measure of SDC severity (i.e. total parcel SDC).

For each patient, we defined measures of FC connector and FC hub damage as weighted means of the control-derived PC and WMD values for damaged parcels as in the study by Gratton et al., (**Fig. 3A**). Replicating the connector damage effect, a 2-predictor multiple regression model identified a significant effect of connector damage, but not hub damage, on modularity (**Fig. 3B, left**, see also **Fig. S2**). However, a 3-predictor model that included total SDC explained significantly more variance (*F*_1,110_=22.6, *p*<0.001) and featured total SDC as the only significant predictor (**Fig 3.B,** top right). This model was not improved by adding lesion volume, and total SDC remained the only significant predictor after lesion volume was added (**Fig. 3B,** bottom). Correlational analyses indicated that modularity was more strongly related to total SDC than to connector damage even when adjusting for lesion volume (**Fig. S2**). These results show that the loss of modularity is better predicted by SDC severity than by damage to regions with critical FC attributes or lesion size.

**Figure 3.**
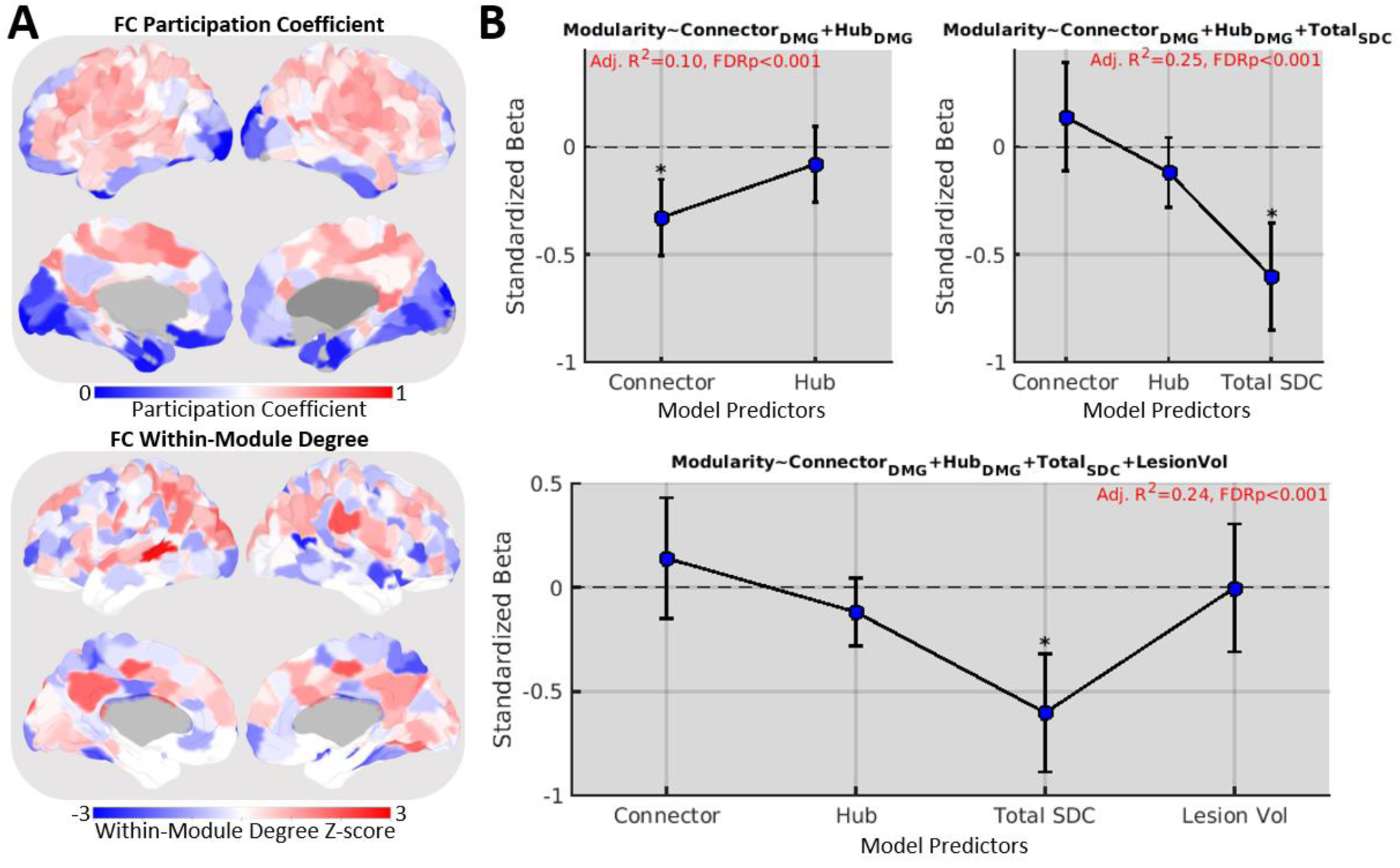
Effects of disconnection and connector node damage on modularity. **A.** FC participation coefficients (top) and within-module degree z-scores (bottom) for each cortical parcel computed based on the mean control FC matrix. **B.** Standardized Betas and 95% confidence intervals for three nested regression models of FC modularity. *FDR*p*<0.05. See also **Figure S2.**

### Disconnection patterns explain functional connectivity disruptions

We next compared the ability of multivariate damage and SDC information to explain FC disruptions. For each FC measure (see **Fig. 2D**), we fit four separate partial least squares regression (PLSR) models using the different structural measures (see **Fig. 1A**) as predictors. The optimal number of PLS components for each model was determined via jackknife cross-validation. To identify the best model of each FC measure, we compared Akaike Information Criterion (AIC) weights among models. AIC weights can be interpreted as conditional probabilities that a model is the best of a set given the data and the set of models (Wagenmakers and Farrel, 2004).

SDC models consistently explained more variance in the FC measures than damage models (**Fig. 4A**). While SDC models tended to use more PLS components than damage models (**Fig. 4B**), subsequent comparisons using AIC weights accounted for differences in model complexity (see also **Fig. S3B**). Together, the parcel and tract SDC models had highest AIC weights for all FC measures (**Fig. 4C**). Parcel SDC models had the highest AIC weights for nine FC measures that included core measures of interhemispheric integration, ipsilesional segregation, and system modularity (red boxes in **Fig. 4**). Thus, SDC measures (particularly parcel SDC) outperformed damage measures for explaining FC disruptions.

**Figure 4.**
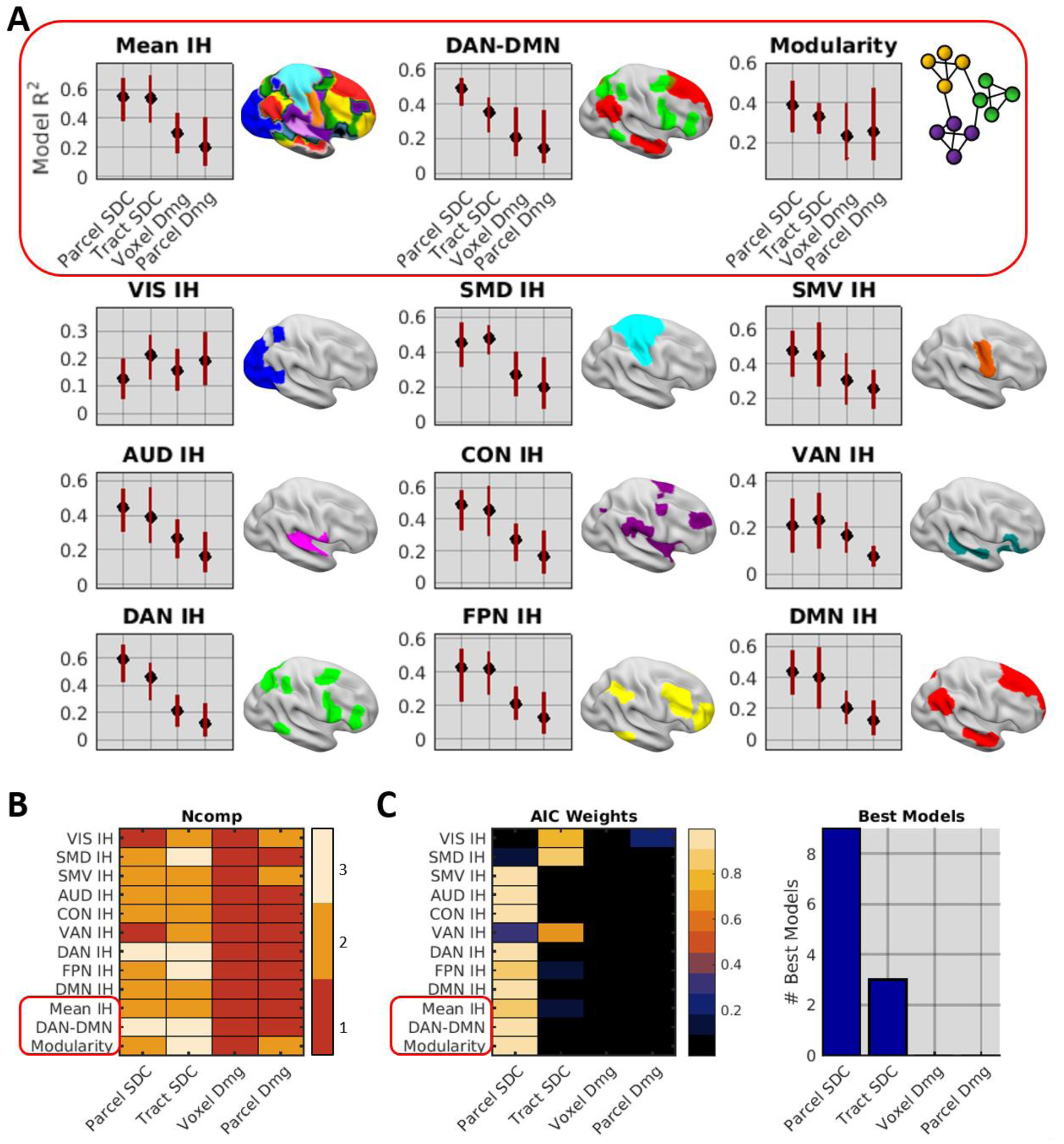
Disconnection explains FC disruptions. **A.** PLSR model fits and FWE-corrected 95% CIs. The red box highlights distinct core FC disruptions. **B.** Number of components included in each model. **C.** (Left) Model AIC weights. (Right) Number of times each model was selected as the best model of the set. See also **Figure S3A-D**.

### Disconnection patterns associated with core FC disruptions

Prior work indicates that disruptions of FC involving different systems are correlated across patients (Baldassarre et al., 2014; Siegel et al., 2016). We observed moderate-to-strong correlations among the different FC measures (bottom triangle in **Fig. 5A**) that were reflected by correlations of the unthresholded PLSR weights from the parcel SDC models (top triangle in **Fig. 5A**). Consistent with the proposal that correlations among FC disruptions reflect the simultaneous disruption of proximal WM pathways within vascular territories (Corbetta et al., 2018), FC disruptions and corresponding parcel SDC weight patterns were highly correlated among systems with dense connections traversing the middle cerebral artery (MCA) territory (**Fig. 5A** – systems other than VIS; see **Fig. S1D**) and weakly-to-negatively correlated between these systems and other systems (**Fig. 5A** – VIS).

**Figure 5.**
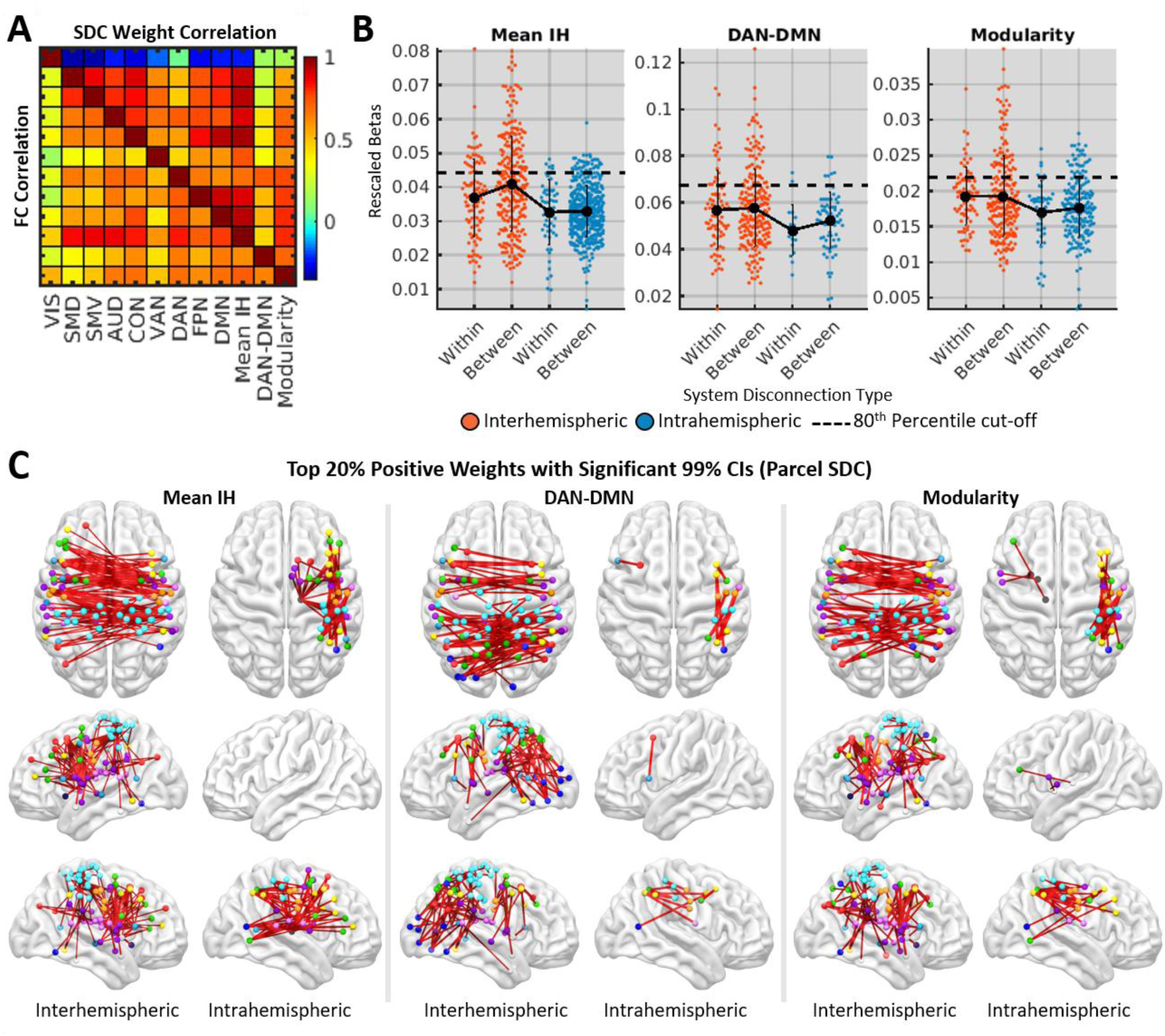
SDC topographies of core FC disruptions. **A.** The bottom triangle shows correlations among the different FC measures. The top triangle shows correlations among the PLSR weight vectors from the corresponding parcel SDC models. **B.** Distributions of significant parcel SDC weights that predicted reduced interhemispheric FC, increased DAN-DMN FC, and reduced modularity. Weights are shown separately for different system and hemispheric connection types. Dashed lines correspond to 80^th^ percentile cut-offs of thresholded weights. Means/SDs are shown as line plots and error bars. **C.** Brain plots show top 20% of significant positive parcel SDC weights (i.e. weights above dashed lines in A). See also **Figures S4-S5**.

We extracted the parcel SDC PLSR model weights with significant 99% CIs (1000 bootstraps) and positive signs (signs were flipped so that positive weights predicted more severe FC disruptions in all models) from the models of mean interhemispheric within-system FC, ipsilesional DAN-DMN FC, and modularity to characterize the most salient SDCs. On average, parcel SDC weights were larger for interhemispheric SDCs than for intrahemispheric SDCs (**Fig. 5B**, compare orange and blue dots), and the top weights for each model corresponded overwhelmingly to interhemispheric SDCs within the MCA territory (**Fig. 5B**, dots above the dashed lines; **Fig. 5C**; see also **Fig. S4**). The top weights also included several intrahemispheric SDCs that primarily corresponded to right thalamocortical, fronto-parietal/fronto-temporal, and/or left frontal SDCs (**Fig. 5C**; see also **Fig. S4**). SDCs involving the cerebellum and brainstem (and to a lesser extent, VIS) were associated with less severe FC disruptions (**Fig. S4-S5**), consistent with the interpretation that the correlation among FC disruptions reflects co-occurring SDCs of diverse commissural and association pathways by MCA stroke.

### A low-dimensional relationship linking disconnection and functional connectivity

The results reported above support the conclusion that SDCs underlie previously identified FC disruptions. However, the core FC disruptions identified by previous studies may not be the only or even the primary consequences of SDCs. To thoroughly characterize the broader relationships between SDC and FC across patients, we applied a partial least squares correlation (PLSC) analysis to the dense parcel-wise FC and SDC data from the patient sample. PLSC identifies latent variables (LVs) that maximally account for the covariance between two datasets by performing a linear decomposition of the cross-block covariance matrix, and produces loadings, scores, and magnitudes (i.e. covariance explained) for each LV.

As noted in the Introduction, we expected a low-dimensional relationship between SDC and FC patterns. Consistent with this interpretation, the first ten LVs identified by the PLSC explained 87% of the covariance between the dense SDC and FC data (**Fig. 6A**). Permutation testing (1000 permutations) revealed that the first two LVs (i.e. LV1 and LV2) each explained significantly more covariance (45% and 21%, respectively) than expected under the empirical null (FDR*p*’s=0.005).

**Figure 6.**
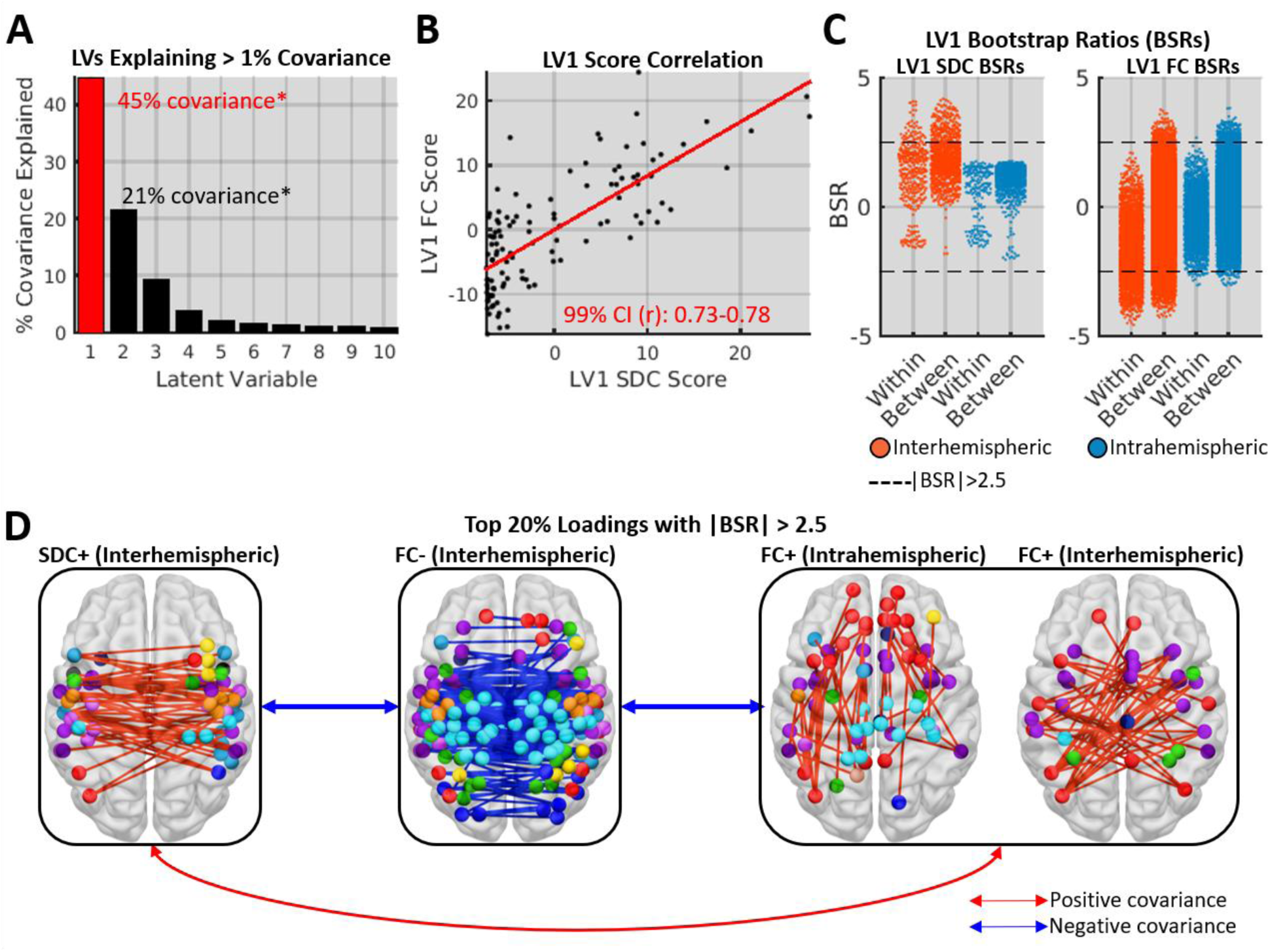
PLSC results. **A.** Covariance explained by 1^st^ 10 LVs. LV1 (red outline) is characterized in subsequent panels. **B.** Relationship between expression of LV1 SDC and FC patterns across patients with bootstrapped CI. **C.** BSR distributions for LV1 SDC and FC loadings by connection type category. Dashed lines denote significance thresholds. **D.** Structure-function relationship captured by LV1. Brain images show top 20% of significant loadings with positive and negative signs. Arrows indicate directions of covariance between structural and functional patterns in LV1. See also **Figures S3E-I, Figure S6**.

For each significant LV, we identified significant loadings (i.e. edges) as indicated by absolute bootstrap (1000 bootstraps) signal-to-noise ratios (BSRs) greater than 2.5. While both LV1 and LV2 accounted for a significant portion of the total covariance, the loadings on LV2 did not achieve significance (**Fig. S6**). We thus focused on characterizing LV1. Mean LV1 SDC (*t*_112_=-1.0, *p*=0.34) and FC scores (*t*_112_=-0.62, *p*=0.53) did not differ significantly between patients with left vs. right hemispheric lesions, indicating that LV1 was similarly expressed in both patient groups.

The relationship between the expression of the SDC and FC patterns captured by LV1 is shown in **Figure 6B**, the LV1 BSR distributions for different connection categories are shown in **Figure 6C,** and the topography of the structure-function relationship captured by LV1 is summarized in **Figure 6D**. Significant SDC loadings had only positive signs (**Fig. 6C-D**, LV1 SDC) and corresponded to interhemispheric cortico-cortical SDCs within and between multiple cortical systems. Significant FC loadings had both positive and negative signs, and loadings with positive signs (**Fig. 6C**, LV1 FC; **Fig. 6D**, right panels) included both interhemispheric and intrahemispheric functional connections but were almost exclusively between different cortical systems, while loadings with negative signs (**Fig. 6C**, LV1 FC; **Fig. 6D**, middle panel) corresponded overwhelmingly to interhemispheric functional connections within and between cortical systems. The most stable negative loadings corresponded to within-system functional connections (**Fig. 6C**, negative values for LV1 FC; top loadings in **Fig. 6D**). Therefore, greater expression of the inter-hemispheric SDC+ pattern was associated with stronger FC between systems (i.e. reducing segregation) and weaker interhemispheric FC especially within-systems (i.e. reducing integration).

### Disconnection-linked functional connectivity patterns are behaviorally relevant

The FC patterns associated with LV1 resembled two core FC disruptions: reductions of within-system interhemispheric FC and increases in between-system FC (**Fig. 6C-D**), suggesting that FC disruptions identified based on behavioral relationships can be interpreted as primary consequences of SDCs. We confirmed this correspondence by correlating the patient scores for LV1 with the *a priori* FC measures (**Fig. 7A**, top row). Patient FC scores for LV1 showed an extremely strong negative correlation with mean interhemispheric within-system FC and moderately strong correlations with the other measures. Similar, but weaker, relationships were observed for LV1 SDC scores (**Fig. 7A**, bottom row). To confirm that LV1 captured a behaviorally relevant structure-function relationship, we correlated LV1 scores with performance scores from language, attention, visual memory, spatial memory, and motor domains. SDC and FC scores showed significant correlations with behavioral impairments in multiple domains (**Fig. 7B**, left), even when adjusting for lesion volume (**Fig. 7B**, right). Thus, LV1 was behaviorally relevant despite being identified based on its ability to account for SDC-FC covariance.

**Figure 7.**
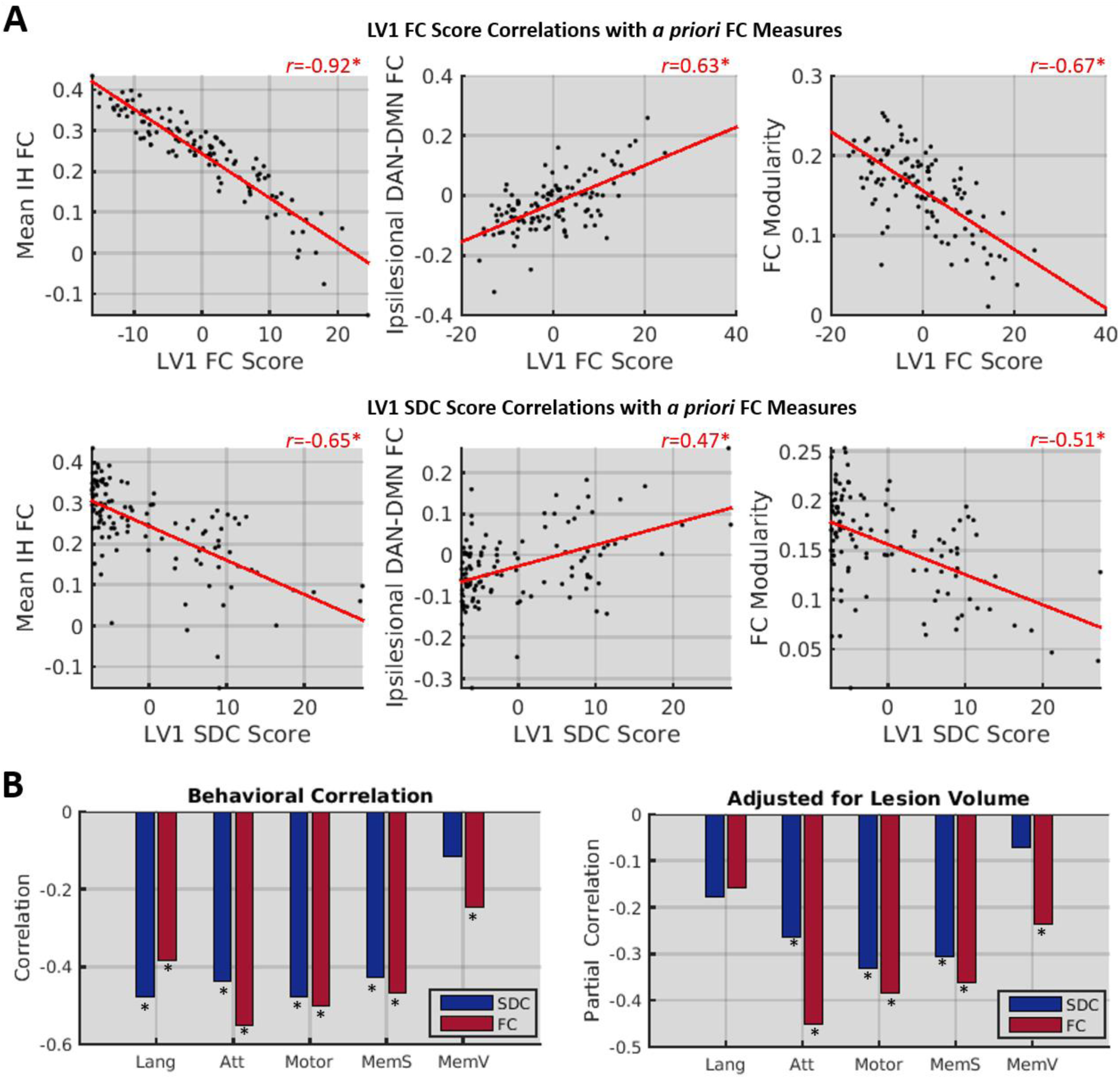
Relationships of LV1 to FC disruptions and behavior. **A.** Scatterplots show relationships between core FC measures and LV1 scores. **B.** Bar plots show correlations between behavioral measures and LV1 scores. Lang – Language, Att – Attention, MemV – Verbal Memory, MemS – Spatial Memory. *FDR*p*<0.05

### Disconnection topographies are partially reflected in functional connectivity patterns

The SC and FC patterns that covary across healthy individuals feature largely divergent topographies, suggesting that they largely reflect indirect network-level relationships (Mišić et al., 2016). However, normal inter-individual variability in SC measures might conceivably be dominated by relatively minor variations around a conserved macroscale architecture that underlies stable group-level FC phenomena like resting-state networks (Greicius et al., 2009; Van Den Heuvel et al., 2009; Shen et al., 2015a). Because SDCs are direct perturbations of this core architecture, we expected that their topographies would be reflected in linked FC patterns.

We assessed the topographic similarity of the SDC and FC components of LV1 by correlating the unthresholded loadings (**Fig. 8A**) common to both components. This analysis revealed a weak but significant negative correlation at the level of the full loading vectors (**Fig. 8B**). Thus, larger positive SDC loadings were weakly associated with larger negative FC loadings for a given edge. Because the degree of structure-function correspondence might be expected to differ among edges with distinct system and/or hemispheric attributes, we separately computed the correlations between the SDC and FC loadings for different hemispheric and system connection categories. **Figure 8C** shows that while considerable correspondence was observed for within-system connections, very little was observed for between-system connections. Supplemental analyses of data from patients with minimal-to-no cortical damage revealed highly consistent results for interhemispheric within-system connections, indicating that this correspondence is not likely to be an artifact of parcel damage (**Fig. S7**). These results support the conclusion that direct perturbations of the structural connectome are directly reflected by changes in the FC of disconnected nodes, particularly for within-system SDCs.

**Figure 8.**
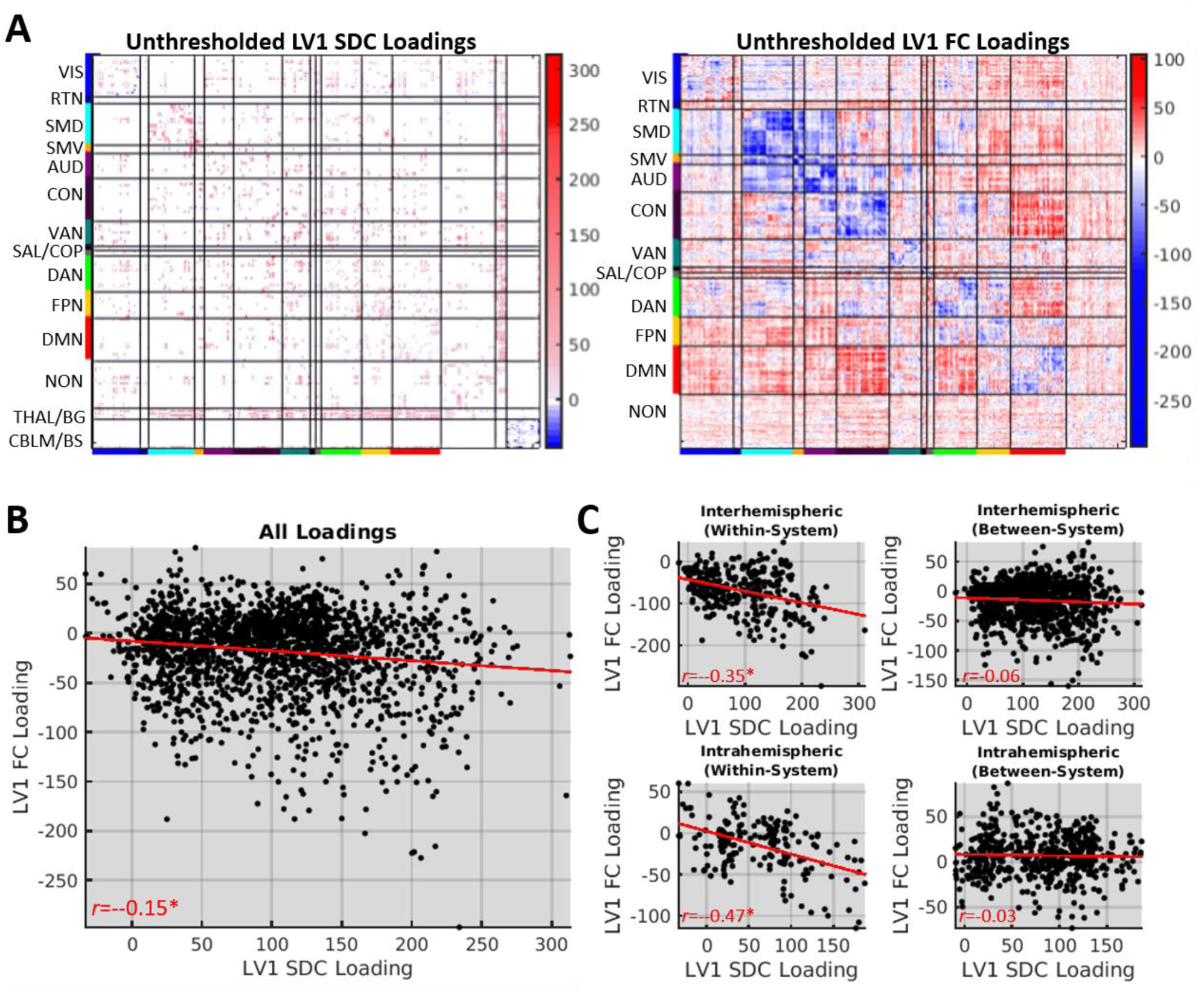
Topographic similarity of LV1 SDC and FC loadings. **A.** Unthresholded LV1 loadings. **B.** Relationship between common LV1 SDC and FC loadings. **C.** Relationships for different connection type categories. *FDR*p*<0.05. See also **Figure S7.**

## Discussion

Stroke disrupts the macroscale functional connectome (Baldassarre et al., 2016a; Carrera and Tononi, 2014; Corbetta et al., 2018; Fox, 2018; Grefkes and Fink, 2014), and a complete understanding of stroke pathophysiology must account for how focal lesions disrupt FC. Our results highlight a key role of SDC in determining the severity of FC disruptions caused by stroke. Specifically, they show that core signatures of stroke are better explained by SDC measures than by focal damage measures. They also reveal a low-dimensional relationship between SDC and FC that is dominated by a single axis linking interhemispheric disconnections to widespread disruptions of FC.

### Structural correlates of functional connectivity disruptions

We tested the hypothesis that SDC is the primary structural factor underlying FC disruptions caused by stroke. Our analyses consistently identified SDC measures as superior to focal damage measures for explaining FC disruptions (**Fig.3-4, Fig. S2-S3**). While parcel SDC measures were generally the most informative, the comparable performance of the tract SDC measures is noteworthy given their relative simplicity (i.e. 70 tracts), as this highlights the explanatory power of even simple SDC measures. Voxel damage models, which included information about WM damage, also outperformed GM-based parcel models (**Fig. 4A; Fig. S5**). Characterization of both the PLSR model weights (**Fig. 5**) and PLSC LV1 loadings (**Fig. 6**) highlighted interhemispheric SDCs as particularly important for explaining FC disruptions, an observation that we will discuss in detail shortly.

Our results argue against a popular account of FC disruptions in patients with focal brain lesions, namely that most focal lesions selectively disrupt function within damaged functional systems (Nomura et al., 2010; Warren et al., 2014) while lesions affecting FC connector nodes broadly disrupt system organization (Gratton et al., 2012). This account has been highly influential, but it has been challenged by recent evidence indicating that lesions can disrupt FC between undamaged systems while sparing FC within damaged systems (Eldaief et al., 2016), that damage to “connector” regions may not be particularly important for predicting FC disruptions in multivariate GM damage models (Yuan et al., 2017), and that disruptions of task-evoked systems are linked to lesions that minimally overlap with constituent regions but that broadly disrupt within-system SC (Griffis et al., 2017a). While we directly replicated the effect of FC connector damage on modularity, adding total parcel SDC information increased the variance explained by a factor of 2.5, and the effect of connector damage did not persist when SDC information was simultaneously modeled (**Fig. 3B**). Direct comparisons also revealed that the effect of connector damage was significantly weaker than the effect of total parcel SDC (**Fig. S2**). More broadly, our PLSR results strongly argue against GM damage as a primary source of FC disruptions in patients with focal brain lesions, as the parcel damage models consistently demonstrated the poorest performance of all models tested (**Fig. 3; Fig. S3A-D**).

Our results are consistent with a general prediction of previous computational studies modeling lesion-induced FC disruptions – namely that a lesion’s impact on FC is strongly influenced by its expected impact on the structural connectome (Alstott et al., 2009; Cabral et al., 2012; Saenger et al., 2017; Váša et al., 2015). We note, however, that a drawback of most computational studies is that they have typically simulated SDCs by node-wise edge deletion for damaged nodes, and this approach cannot account for the effects of WM damage, which is common (**Fig. 1**). In contrast, our approach to measuring expected SDCs does account for the effects of WM damage. The incorporation of similar approaches into future modeling studies might enable more realistic simulations and thus improve the generalizability of specific findings.

Our results also complement prior studies that have implicated WM damage and SDC as important factors in determining the severity of cognitive and behavioral deficits after stroke (Catani et al., 2012; Chechlacz et al., 2013; Corbetta et al., 2015; Forkel et al., 2014; Griffis et al., 2017b, 2017a, Kuceyeski et al., 2015, 2016a; Marebwa et al., 2017; Thiebaut De Schotten et al., 2014; Yourganov et al., 2016). Speculatively, these relationships may be partially mediated by the large-scale functional disruptions precipitated by SDCs. Future studies using path modeling or mediation analyses are an important next step towards understanding the complete relationship linking focal lesions to functional disruptions and behavioral impairments.

### Low-dimensional covariance between disconnection and functional connectivity

We empirically characterized how perturbations of the structural connectome are reflected in whole-brain FC patterns. Our analyses revealed a low-dimensional relationship between SDC and FC, such that two-thirds of the SDC-FC covariance could be attributed to two LVs (**Fig. 6A**). Principal component analyses (PCAs) indicated that this result did not simply reflect an intrinsic low dimensionality of the lesion, SDC, or FC data (**Fig. S8**).

This analysis was partially motivated by the consideration that the FC patterns that maximally covary with SDCs might be distinct from the core FC disruptions reported in the literature. However, the FC pattern captured by LV1 clearly reflected core FC disruptions (**Fig. 6; Fig. 7A**) that have been identified based on their relationships to behavior (Baldassarre et al., 2014, 2016b, 2018; Carter et al., 2010; He et al., 2007; Tang et al., 2016) and deviations from controls (Ramsey et al., 2016; Siegel et al., 2016). Accordingly, patient-level expression of LV1 correlated with behavioral impairments (**Fig. 7B**). This was particularly pronounced for attention, spatial memory, and motor domains, consistent with the notion that cortico-cortical FC is critical for higher cognitive functions (Corbetta et al., 2018; Siegel et al., 2016) and with previous reports of corrrelated motor and attention deficits (Baldassarre et al., 2016). Because the FC profile associated with LV1 was identified based on its relationship to SDCs rather than behavior, we speculate that it represents an aggregate of different behaviorally relevant FC topographies that covary with interhemispheric SDCs. Consistent with this interpretation, the LV1 FC pattern qualitatively resembles the FC pattern that was found to predict multi-domain behavioral deficits in a previous analysis of FC data from this sample (Siegel et al., 2016).

The relationships shown in **Figure 7B** are consistent with evidence indicating that behaviorally relevant FC abnormalities are correlated across patients and tend to vary in topography and severity but not form (Corbetta et al., 2018). The weakening of interhemispheric FC is a common signature of impairments in multiple behavioral domains, and domain-specific deficits are associated with specific topographies of weakened connections (Baldassarre et al., 2014, 2016b, 2018; Bauer et al., 2014; Carter et al., 2010; He et al., 2007; Siegel et al., 2016; Tang et al., 2016). Our results advance a mechanistic explanation for the low-dimensionality of behavioral and FC disruptions after stroke. Specifically, they suggest that strokes within the MCA territory disrupt interhemispheric SC both within and between cortical systems, producing both direct and network-level effects on FC that lead to a breakdown in the balance of system integration and segregation.

Why might interhemispheric SDCs be particularly important for determining the functional consequences of focal brain lesions? One putative explanation is that stable interhemispheric integration is a fundamental component of large-scale FC organization (Shen et al., 2015a,b) that shapes broader aspects of FC. This explanation is broadly consistent with computational modeling work indicating that the interaction between callosal SC and physiological factors may play an important role in mediating FC between other regions, including those that lack direct SC (Messé et al., 2014), and with reports indicating that interhemispheric SDCs both reduce interhemispheric FC and increase intrahemispheric FC in non-human primates (O’Reilly et al., 2013). Nonetheless, this is an open question that should be addressed in detail by future work.

The observed low-dimensional relationship between SDC and FC contrasts sharply with recent reports of high-dimensional covariance between SC and FC patterns in healthy individuals. For example, a recent PLSC study reported that 5 LVs explained only 22.6% of the covariance between SC and FC patterns in a sample of 156 healthy participants (Mišić et al., 2016). In contrast, the first 5 LVs identified by our PLSC analysis together explained 82% of the structure-function covariance across patients (**Fig. 6A**). One explanation for this stark difference is that normal variability in MRI measures of SC reflects relatively minor but not inconsequential variations around an otherwise conserved structural scaffold that shapes the core FC topographies identified by group-level analyses, while variability in SDC measures reflects major differences in the integrity of this core scaffold. This explanation is consistent with the fact that the topographies of the SC and FC components of the LVs identified by Misic et al., were discordant, while the SDC and FC components of LV1 identified by our analysis exhibited significant topographic similarity (**Fig. 8**). Our data show that direct perturbances of the structural connectome produce changes in FC that partially mirror the affected structural connections, and these effects may be most consistent and reliable for interhemispheric within-network SDCs (**Fig. 8**).

Even so, the correspondence between the linked SDC and FC patterns was still relatively weak (**Fig. 8**), and many of the most stable SDC loadings corresponded to connection types that showed weak topographic similarity (e.g. see interhemispheric between-system in **Fig. 6C, Fig. 8C**). This suggests that much of the covariance between SDC and FC may reflect indirect SDC effects that arise via propagation along serial connections in polysynaptic pathways (Carter et al., 2012; Lu et al., 2011) or through larger-scale network dynamics (Adachi et al., 2011; Alstott et al., 2009; Mišić et al., 2016) arising from the sudden loss of diverse afferent and/or efferent connections throughout distributed brain systems. Future studies should aim to characterize the nature of putative indirect SDC effects.

### Considerations for studying structure-function relationships in the lesioned brain

The current study featured several methodological advantages over previous work on this topic. First, we explicitly incorporated SDC measures. This increased our ability to account for the effects of interest beyond what would have been possible using focal damage measures alone. Previous studies have typically used parcel damage and related approaches that focus on the GM to study lesion-FC relationships (Gratton et al., 2012; Nomura et al., 2010; Ovadia-Caro et al., 2013; Yuan et al., 2017). This both ignores potentially relevant information about WM damage (e.g. compare voxel and parcel damage model fits in **Fig. 4**) and may lead to the mis-localization of WM effects into nearby GM parcels (**Fig. S5**). This might occur when the most relevant variables (in this case, WM voxels) are not measured, but proxy information is available from correlated measured variables (i.e. GM parcels). Hypothetically, such effects could also lead to distorted SDC weight topographies if outcomes were largely driven by GM damage that correlated with specific SDCs, and we accounted for this possibility by performing an additional PLSR analysis that included both parcel damage and parcel SDC measures as predictors (see Methods). This had minimal effect on parcel SDC weight topographies but substantially altered the parcel damage weight topographies. While this was expected given that SDCs provided the most relevant information about FC, it emphasizes the importance of accounting for potentially relevant lesion effects when drawing conclusions about critical locations or SDC topographies in lesion analyses.

We also used data collected from a relatively large sample of first-ever stroke patients at the sub-acute stage of recovery, while most previous studies utilized data obtained from relatively small samples (i.e. *n*=12-35) of patients (Eldaief et al., 2016; Gratton et al., 2012; Nomura et al., 2010; Ovadia-Caro et al., 2013) with diverse lesion etiologies (e.g. stroke, traumatic brain injury, tumor resection, etc.) at varying stages of recovery (Eldaief et al., 2016; Gratton et al., 2012; Nomura et al., 2010; Yuan et al., 2017). Larger patient samples increase power and stabilize effect estimates (Poldrack, 2012; Yarkoni, 2009), but the importance of studying patients with similar lesion etiologies at the same phase of recovery warrants further discussion.

All lesions are not created equal – the pathological origin of a given lesion contributes to its physiological and behavioral consequences (Anderson et al., 1990). For example, slow-growing lesions (e.g. tumors) tend to produce less severe cognitive and behavioral deficits than sudden-onset lesions (e.g. ischemic stroke). This may reflect differences in the brain’s ability to compensate for lesions that develop on different time-scales (Desmurget et al., 2007). Lesions resulting from specific pathologies may also exhibit spatial biases - while medial prefrontal regions are frequently affected by low-grade glioma (Duffau and Capelle, 2004), they are infrequently affected by stroke (Corbetta et al., 2015; Sperber and Karnath, 2016). Further, the behavioral and physiological consequences of brain lesions are not static but evolve throughout the course of recovery (Corbetta et al., 2005; Heiss et al., 1999; Ramsey et al., 2016; Rehme et al., 2011; Saur et al., 2006; Siegel et al., 2018). Therefore, it is ideal to utilize data from patients with similar lesion etiologies and in the same recovery phase to minimize the risk of latent confounds in lesion analyses.

### Limitations

SDC measures were defined for each patient based on the intersection of their lesion with a structural connectome atlas constructed with data from healthy individuals (see Methods). Similar atlas-based approaches have been used by other recent lesion studies (Carter et al., 2017; Foulon et al., 2018a; Griffis et al., 2017b, 2017a; Hope et al., 2018; Kuceyeski et al., 2013, 2014, 2015, 2016b; Pustina et al., 2017a; Ramsey et al., 2017), and analogous strategies are often employed to study SC-FC relationships in animal models (Adachi et al., 2011; Grandjean et al., 2017; Grayson et al., 2016; Shen et al., 2015b). While these approaches assume similar approximations of individual structural connectomes by the atlas and cannot account for interindividual variability in the properties of un-damaged fiber pathways (Forkel and Catani, 2018; Forkel et al., 2014), they also offer some protection against potential biases arising from inter-individual differences in diffusion MRI data quality, lesion effects on data processing/reconstruction, etc. They also provide an intuitive means of estimating the severity of SDC relative to a common reference (i.e. % of template streamlines affected) that can be employed consistently across samples and/or studies. Further, the structural connectome atlas used in the current study was based on very high-quality data (i.e. 90 direction high-angular resolution diffusion imaging) from a very large sample of participants (i.e. N=842) and was expert-vetted to reduce the likelihood of false positive connections (Yeh et al., 2018). While direct patient SDC measures would be ideal, our results nonetheless show that the atlas-based measures provided important information about FC beyond what is present in focal damage measures.

A similar limitation is that template-based areal parcellations may not provide comparable approximations of areal boundaries for all participants. Previous analyses of data from this sample have shown that the parcellation delineates largely homogenous functional parcels in both patients and controls (Siegel et al., 2018), but inter-individual variability in areal boundaries and/or system topographies could still influence our measures (Braga and Buckner, 2017; Gordon et al., 2017; Gratton et al., 2018; Marek et al., 2018). However, template-based parcellation approaches have advantages that are analogous to those described for template-based disconnection approaches. Further, the large amount of necessary data (Gordon et al., 2017) and the potential for distortions by lesion and/or hemodynamic factors (Siegel et al., 2017) make individual functional parcellations currently infeasible in sub-acute stroke patients.

Previous studies have successfully used measures of expected SDC to model behavioral impairments (Foulon et al., 2018b; Fridriksson et al., 2013; Griffis et al., 2017b; Kuceyeski et al., 2015, 2016b) and changes in brain structure (Foulon et al., 2018b; Kuceyeski et al., 2014), but SDC information may not always provide unique information beyond what is present in focal damage measures (Hope et al., 2018). We expect that the utility of SDC information will likely depend on several factors that include the degree to which the variable of interest depends on SDC vs. focal damage, the quality of the SDC measures, and the lesion characteristics of the patient sample. Because SDC information is implicit in the lesion, it is likely that voxel-based lesion measures will provide similar information as SDC measures when lesion coverage and diversity is sufficiently high to recover the implicit SDC information, although this would likely require huge samples with diverse lesions and dense coverage throughout the brain (e.g. *N*=818 in Hope et al., 2018). Even in scenarios where SDC and damage information enable similar prediction, we consider the inclusion of SDC information useful from a neuroscientific perspective.

## Acknowledgments

Funding: This research was supported by grants RO1 NS095741 and R01 HD061117 to M.C.

Data were provided [in part] by the Human Connectome Project, WU-Minn Consortium (Principal Investigators: David Van Essen and Kamil Ugurbil; 1U54MH091657) funded by the 16 NIH Institutes and Centers that support the NIH Blueprint for Neuroscience Research; and by the McDonnell Center for Systems Neuroscience at Washington University.

We thank Alexandre Carter for assisting with lesion segmentation, and Joshua Siegel for comments on earlier versions of the manuscript.

## Author Contributions

J.G and G.S designed the analyses and wrote the paper. J.G and N.M. performed data processing and analyses. J.G., G.S., and M.C. edited the paper. G.S. and M.C. contributed data and other resources.

## Methods

### Contact for Reagent and Resource Sharing

Requests for additional information or resources should be directed to the Lead Contact, Gordon Shulman.

### Participant information

Patients and controls provided written informed consent prior to participation in the study. Study procedures were performed in accordance with the Declaration of Helsinki ethical principles and approved by the Institutional Review Board at Washington University in St. Louis. The complete data collection protocol is described in detail in our previous publication (Corbetta et al., 2015). Data from 132 first-time stroke patients who presented with clinical evidence of cognitive and/or behavioral impairment and data from 33 demographically matched healthy controls were considered for inclusion in the study. Data from 114 patients and 24 controls met quality control criteria (described below) and were included in the study. Participant demographics are shown in **Table 1.**

### Neuroimaging data collection

Neuroimaging data were collected using a Siemens 3T Tim-Trio scanner at the Washington University School of Medicine with a 12-channel head coil, and are fully described elsewhere (Corbetta et al., 2015; Siegel et al., 2016). Sagittal T1-weighted MP-RAGE (TR=1950 msec; TE=2.26 msec, flip angle = 90 degrees; voxel dimensions = 1.0×1.0×1.0 mm), transverse T2-weighted turbo spin-echo (TR=2500 msec; TE=43 msec; voxel dimensions = 1×1×1), and sagittal T2-weighted FLAIR (TR=750 msec; TE=32 msec; voxel dimensions = 1.5×1.5×1.5 mm) structural scans were obtained along with gradient echo EPI (TR=2000 msec; TE=2 msec; 32 contiguous slices; 4×4 mm in-plane resolution) resting-state functional MRI scans. During the fMRI scans, participants were instructed to fixate on a small white centrally-located fixation cross presented against a black background on a screen at the back of the magnet bore. An Eyelink 1000 eye-tracking system (SR Research) was used to monitor when participant’s eyes were opened/closed during each run. Between six and eight resting-state scans (128 volumes each) were obtained from each participant (∼30 minutes total).

### Lesion identification

Lesions were manually segmented on each patient’s structural MRI scans using the Analyze software package (Robb and Hanson, 1991). Surrounding vasogenic edema was included in the lesion definition for patients with hemorrhagic stroke. All segmentations were reviewed by two board certified neurologists (Maurizio Corbetta and Alexandre Carter), and were reviewed a second time by MC. The final segmentations were used as binary lesion masks for subsequent processing and analysis steps. Lesion masks were transformed to MNI atlas space using a combination of linear transformations and non-linear warps and were resampled to have isotropic voxel resolution.

### Behavioral measures

Participants performed a behavioral battery consisting of multiple assessments within motor, language, attention, verbal memory, spatial memory, and visual domains. Principal components analyses (PCA) were used to decompose the behavioral data from each domain as described in our previous publication (Corbetta et al., 2015). Detailed descriptions of the behavioral testing and PCA analyses can be found in the Supplemental Material for (Corbetta et al., 2015; Siegel et al., 2016). Analogously to other previous work (Ramsey et al., 2017; Siegel et al., 2016, 2018), the first PCs from each behavioral domain (with the exception of vision) were considered as domain scores of interest, and were used in analyses that related imaging measures to behavior (**Fig. 7B**). Of the 114 patients that were included in the main analyses, 108 had data for the language domain, 93 had data for the attention domain, 101 had data for the motor domain, and 84 had data for the verbal and spatial memory domains.

### MRI data processing

Functional MRI data pre-processing consisted of slice-timing correction using sinc interpolation, correction of inter-slice intensity differences resulting from interleaved acquisition, normalization of whole-brain intensity values to a mode of 1000, correction for distortion via synthetic field map estimation, and within-and between-scan spatial re-alignment. BOLD data were re-aligned, co-registered to the corresponding structural images, normalized to atlas space, and resampled to 3mm cubic voxel resolution using a combination of linear transformations and non-linear warps. Prior to estimating FC, additional processing steps were applied to account for non-neural sources of signal variance. Confounds related to head motion, global signal fluctuations, and non-grey matter signal compartments were removed from the data by regression of the six head motion parameters obtained from rigid body correction, along with the global GM signal and the CSF and white matter signals extracted from FreeSurfer tissue segmentations (Dale et al., 1999). BOLD data were band-pass filtered (0.009 < *f* < 0.08 Hz) to retain low-frequency fluctuations. A frame was censored if it exceeded a 0.5 mm framewise displacement threshold, and the succeeding frame was also censored to further reduce confounds related to motion (Power et al., 2014). The first four frames of each run were discarded to allow for the scanner to achieve steady-state magnetization.

Cortical surface generation and subsequent fMRI data processing generally followed previously published minimal preprocessing procedures (Glasser et al., 2013), although some modifications were required to accommodate lesioned brains (Siegel et al., 2016, 2017). FreeSurfer was used to automatically obtain anatomical surfaces from the T1-weighted structural scans (Dale et al., 1999; Fischl et al., 1999), and the resulting segmentations were visually inspected to ensure accuracy. Data from patients with failed registrations and/or segmentations were modified by replacing the values of lesioned voxels with normal values from the structural atlas prior to running the registration and segmentation procedures, and the modified voxels were masked out after running the procedures (Siegel et al., 2017). Each hemisphere was resampled to 164,000 vertices, and the two hemispheres were registered to each other (Van Essen et al., 2001). The data were then down-sampled to 32,000 vertices per hemisphere. Ribbon-constrained sampling in Connectome Workbench was used to sample functional MRI volumes to each participant’s individual surface, and voxels with coefficients of variation > 0.5 standard deviations above the mean of all voxels within a 5 mm sigma Gaussian neighborhood were excluded from volume to surface mapping (Glasser et al., 2013).

Participants were excluded if they had less than 180 usable frames of resting-state data after applying quality controls, and this resulted in the exclusion of 18 patients and 9 controls. The remaining 114 patients and 24 controls who had sufficient FC data were included in the primary analyses.

### Parcels and system assignments

We used the Gordon333 cortical parcellation and system community assignments (available at http://www.nil.wustl.edu/labs/petersen/Resources.html) to obtain parcel-level and system-level measures of functional connectivity. This parcellation is based on functional connectivity boundary mapping and InfoMap community detection analyses of resting-state fMRI data from 120 healthy individuals (Gordon et al., 2016), and consists of 333 cortical parcels associated with 13 large-scale systems. Previous studies involving the current dataset excluded 9 parcels for having very low numbers of vertices, and so they were also excluded here for consistency (Siegel et al., 2016, 2018). The remaining 324 surface-based cortical parcels were used for subsequent surface-based estimation of functional connectivity.

In addition to the 324 cortical parcels, we also defined a set of 35 sub-cortical and cerebellar parcels to allow for the complete quantification of damage and disconnection throughout the brain. This set of parcels consisted of 34 parcels from the automatic anatomical labeling (AAL) atlas (Tzourio-Mazoyer et al., 2002) that corresponded to different portions of the thalamus, basal ganglia, and cerebellum, and also included 1 parcel from the Harvard-Oxford Subcortical Atlas that corresponded to the brainstem (https://fsl.fmrib.ox.ac.uk/fsl/fslwiki/Atlases). The full set of 359 parcels is shown in **Figure S1** and was used for subsequent estimation of regional damage and structural disconnection. Cortical parcel borders were removed as they sometimes contained values that were inconsistent with the adjacent parcels. The borderless cortical parcels were dilated by 2mm to improve the sensitivity of subsequent structural connectivity analyses (Van Den Heuvel et al., 2009; Wilson et al., 2011) and to allow for a slightly relaxed threshold for determining cortical damage (Pustina et al., 2017b).

### Functional connectivity estimation

Parcel-wise FC matrices were constructed by correlating the average (i.e. across all within-parcel vertices) nuisance-regressed BOLD timeseries of each surface parcel with the average nuisance-regressed BOLD timeseries of every other parcel and applying the Fisher z-transformation to the resulting linear correlation values. For each patient, vertices that fell within the boundaries of the lesion were masked out, and parcels with less than 60 vertices remaining after excluding lesioned vertices were completely excluded by setting the values to NaN, analogously to previous reports (Siegel et al., 2016; Siegel et al., 2018). We note that analyses that were performed without removing lesioned vertices produced very similar results for all analyses (not shown), likely owing to the relatively low frequency of cortical lesions in our sample (**Fig. 1B, Fig. S7A**).

### Structural connectome template

We used a publicly available tractography atlas (available at www.brain.labsolver.org) to create a structural connectome template. For each patient, we then estimated structural disconnections by intersecting the template with the lesion (**Fig. 1A**). The tractography atlas was constructed using data from 842 Human Connectome Project participants (Yeh et al., 2018), and atlas data were accessed under the WU-Minn HCP open access data use term. The full details regarding the construction of the tractography atlas can be found elsewhere (Yeh et al., 2018). To summarize, Yeh and colleagues (2018) reconstructed the high-angular resolution diffusion MRI data (b-values: 1000, 2000, and 3000 s/mm2; diffusion sampling directions: 90, 90, and 90; in-plane resolution: 1.25mm) from 842 Human Connectome Project participants in the MNI template space using Q-space diffeomorphic reconstruction (Yeh and Tseng, 2011), averaged the resulting spin distribution functions (SDFs) to obtain population-level streamline trajectories (available as the HCP-842 template at www.dsi-studio.labsolver.org), and performed deterministic fiber tracking (Yeh et al., 2013b) to extract 550,000 streamline trajectories that were then vetted and labeled by a team of neuroanatomists (for detailed descriptions of the procedures, see Yeh et al., 2018). Thus, the tractography atlas consisted of expert-vetted end-to-end streamline trajectories in the MNI template space that were each associated with 1 of 66 neuroanatomically defined fiber bundles (e.g. superior longitudinal fasciculus, corpus callosum, etc.) that are grouped into commissural, association, projection, brainstem, and cerebellar pathways (cranial nerves were not included). Because we expected that different segments of the corpus callosum might show different relationships to FC in the tract disconnection analyses, we split the corpus callosum into 5 segments based on the FreesurferSeg ROIs included with DSI_studio.

We used command line utilities provided in the DSI_studio software package (available at www.dsi-studio.labsolver.org) to define the normative parcel-wise structural connectome template based on the tractography atlas (**Fig. S1**). To define the parcel-wise structural connectome, we first combined the labeled streamline bundles from the structural connectome atlas (e.g. short-range U-fibers, callosal projections, etc.) into a single aggregate *.trk* file, and then extracted all streamlines that bilaterally terminated (i.e. began and ended) within any pair of the 359 volume-based parcels. This resulted in a 359×359 structural connectivity adjacency matrix *A*^*S*^ where each entry *A*^*S*^_*ij*_ indexed the number of streamlines connecting parcel *i* and parcel *j*. Due to the close proximity of ventral visual parcels and dorsal cerebellar parcels, a small number of dorsal cerebellar streamlines were captured by the dilated visual parcels. Therefore, we removed any connections between visual areas and the cerebellum.

### Functional connectivity measures

Functional connectivity matrices were averaged from each group to create group-averaged FC matrices (**Fig. 2B**). The mean matrix from the control group was subtracted from the mean matrix from the patient group to create a difference matrix (**Fig. 2C**). Linear correlations between the upper triangles of the group-averaged and difference matrices were used to assess the similarity of mean FC topographies with each other and with the difference matrix.

We defined 12 *a priori* measures of network dysfunction based on previously reported FC abnormalities in sub-acute stroke patients. For each of nine bilateral functional systems, we averaged the FC strengths over all within-system interhemispheric functional connections. This resulted in nine system-specific interhemispheric FC measures (**Fig. 2D**, left). To obtain a general measure of interhemispheric integration, we averaged the nine system-specific interhemispheric FC measures to obtain a single measure of mean interhemispheric within-system FC. We note that this measure essentially corresponded to the first principal component of the nine interhemispheric FC measures (*R*^*2*^=0.94), which explained 70% of the total variance across the nine system-specific interhemispheric FC measures. To obtain a measure of system segregation, we averaged the FC strengths for all ipsilesional DAN and DMN functional connections (**Fig. 2D**, middle). We chose this measure because previous analyses data from this sample have reliably reported reduced segregation between the DAN and DMN in patients (Ramsey et al., 2016; Siegel et al., 2016). Finally, to obtain a global measure that considered both integration and segregation, we measured FC modularity (Newman’s Q) using the *community_louvain.m* function from the Brain Connectivity Toolbox (Rubinov and Sporns, 2010). Modularity estimation was performed using the 280 parcels with specific *a priori* system assignments (i.e. excluding unassigned parcels), and modules were defined *a priori* as the default parcel system assignments for the reasons described in Siegel et al., (2016). As in previous studies that have measured modularity in patients with focal brain lesions (Gratton et al., 2012; Siegel et al., 2018), we performed our analyses across multiple edge density thresholds ranging from 4% and 20% edge density in 2% steps (**Fig 2D**, right). The final modularity measure was obtained by averaging over edge density thresholds. We considered this appropriate as modularity estimates were highly correlated across thresholds such that single principal component accounting for 95% of the total variance across thresholds (this component was almost perfectly colinear with the mean across thresholds – *R*^2^=0.99). Unequal variance t-tests were used to compare the *a priori* measures between patients and controls, and false discovery rate (FDR) correction was used to correct for multiple testing (Benjamini and Hochberg, 1995). Results of these analyses are shown in **Figure 2D**.

The parcel-level participation coefficients and within-module degree z-scores shown in **Figure 3A** were estimated by applying the Brain Connectivity Toolbox functions *participation_coef* and *module_degree_zscore* to the mean FC matrix from the control group (shown in **Fig. 2B**) using the same parcels and range of edge density thresholds as the modularity analyses (described above), and averaging across thresholds. Note that FC between parcels with Euclidean distances of less than 20mm was not used in the computation of these measures (Power et al., 2013). The FC connector and hub damage measures were defined according to the same procedure described by Gratton et al., (2012). For each patient, this involved multiplying the amount of damage to each parcel by its participation coefficient (FC connector damage) or its within-module degree z-score (FC hub damage) and then averaging measures over parcels. This produced a single FC connector damage measure and a single FC hub damage measure for each patient.

### Structural lesion features

MATLAB scripts utilizing functions from the matlab_nifti toolbox (available at https://www.mathworks.com/matlabcentral/fileexchange/8797-tools-for-nifti-and-analyze-image) were used to obtain voxel-wise damage and parcel-wise grey matter damage measures. For each patient, we re-shaped their 3×3×3mm lesion mask into a 1-dimensional vector indexing the presence vs. absence of damage at each voxel within the group-level lesion coverage area (hereafter referred to as “voxel damage”). We also computed the proportion of each grey matter parcel that overlapped with each patient’s lesion to create a 1-dimensional vector indexing the amount of damage to each parcel within the group-level lesion coverage area (hereafter referred to as “parcel damage”). MATLAB scripts implementing command line functions from the DSI_studio software package (available at www.dsi-studio.labsolver.org) were used to obtain expected disconnections for each patient based on the intersection of their MNI-registered lesion and the structural connectome template.

For each patient, we extracted all streamlines that passed through the lesion to obtain a 359×359 structural disconnection adjacency matrix *A*^*D*^ where each entry *A*^*D*^_*ij*_ indexed the number of streamlines connecting parcel *i* and parcel *j* that intersected the lesion (i.e. that were disconnected in that patient). We then normalized each structural disconnection matrix *A*^*D*^ via element-wise division by the structural connection matrix *A*^*S*^, such that entries in the resulting matrix *A*^*Dnorm*^ indexed the proportion of streamlines connecting parcel *i* and parcel *j* that were disconnected by the lesion. This step accounted for differences in the number of streamlines connecting different parcel pairs and ensured that all disconnection measurements were directly comparable and intuitively interpretable in terms of proportional disconnection rather than number of streamlines. The upper triangles (excluding diagonal elements) of the normalized disconnection matrices were then extracted and reshaped into a 1-dimensional vector indexing the amount of disconnection for each edge (hereafter referred to as “parcel disconnection”). For each patient, we also calculated the proportion of each neuroanatomically defined fiber bundle from the structural connectome atlas that was disconnected by the lesion, resulting in a 1-dimensional vector indexing the amount of disconnection for each tract (hereafter referred to as “tract disconnection”). Prior to performing any statistical analyses, the damage/disconnection vectors from all 114 patients were stacked on top of each other to form four separate data matrices.

Damage and disconnection frequency maps (**Fig. 1**) were created using the voxel-wise damage and parcel-wise disconnection data. Voxel damage maps were summed across patients, resulting in a map that quantified the number of patients with damage to each voxel in the brain. Parcel disconnections were binarized at a 1% disconnection threshold and summed across patients, resulting in a map that quantified the number of patients with damage to each edge in the structural connectome. The binarized parcel SDCs and parcel damage measures were summed across patients to obtain the number of affected patients at each edge/parcel, and the resulting measures were used to create the histograms shown in **Figure 1C**. The binarized parcel SDC and parcel damage measures were also summed across edges to obtain the total number of parcel SDCs and the total number of damaged parcels for each patient, and these total parcel SDC and total parcel damage measures were used for the analyses summarized in **Figure 1D-E** as described in the next section.

### Multiple linear regressions and partial correlations

To assess the extent to which the extent of parcel SDCs reflected the extent of parcel damage, we regressed total parcel SDC on total parcel damage. The resulting R^2^ value quantified the amount of variance shared between the two measures (**Fig. 1D**). To determine whether the total parcel SDC and total parcel damage measures differed in their dependence on lesion volume, we computed linear correlations between each measure and total lesion volume and compared the strength of the correlations using Steiger’s *z*-test (**Fig. 1E**).

We used multiple linear regressions to compare the effects of FC connector damage, FC hub damage, and total SDC on FC modularity (**Fig. 3**). We first fit a linear regression model that included FC connector damage and FC hub damage as predictors. We then added total SDC to the model to determine whether total SDC explained additional variance beyond what could be attributed to FC connector damage and FC hub damage (i.e. *F*-test on *R*^2^ change statistic), and to simultaneously evaluate the effects of total SDC, FC connector damage, and FC hub damage in a single model. We then added lesion volume to determine whether the same effects were observed when lesion volume was included in the model. Effects were considered significant if they survived FDR correction at 0.05.

Because the original study by Gratton et al., (2012) used a correlation (i.e. rather than regression) approach, and because this allowed us to directly compare the strength of the relationships between different structural measures and modularity, we also performed correlational analyses (**Fig. S2**). For these analyses, we first computed the linear correlations between FC modularity and total SDC, FC connector damage, and FC hub damage. We then compared the correlations between FC connector damage and FC hub damage, and between FC connector damage and total SDC using Steiger’s *z*-tests. These analyses were then repeated after adjusting for lesion volume (i.e. partial correlation). Because the comparison between FC connector damage and FC hub damage was intended to replicate the effect reported by Gratton et al., we used a one-tailed test (i.e. Connector damage > Hub damage). Two-tailed tests were used to compare the correlations for total SDC and FC connector damage. Effects were considered significant if they survived FDR correction at 0.05.

### Partial least squares regressions

We used partial least squares regressions (PLSR) to predict our *a priori* FC measures from our structural damage and disconnection measures (**Fig. 4-5**). PLSR is a multivariate regression technique (Wold et al., 2001) that is closely related to principal components regression (PCR) (Hotelling, 1957). Both PLSR and PCR are particularly useful for situations where there are more variables than observations and/or when there is high collinearity amongst the predictor variables. However, PLSR has important advantages over PCR (Abdi and Williams, 2010) that are primarily due to differences in the criteria used for decomposition of the predictor matrix. Namely, while PCR decomposes the predictor matrix **X** into a set of linearly independent components that maximally account for the variance in **X** and uses the scores on some subset of those components to predict **Y**, PLSR performs a dual decomposition of **X** and **Y** to obtain components from **X** that maximally account for the covariance with **Y**. This typically results in simpler models, and is advantageous over PCR because it reduces the potential for important variables to be omitted from the model on the basis that they explain only small amounts of the variance in **X** (Abdi, 2007, 2010; Krishnan et al., 2011). Detailed descriptions of the theory and algorithms behind the PLSR approach can be found elsewhere (Abdi, 2007, 2010; Krishnan et al., 2011; McIntosh and Lobaugh, 2004; Tie Jong, 1993; Wold et al., 2001). We performed PLSR using the SIMPLS algorithm as implemented in the *plsregress* function included with the MATLAB Statistics and Machine Learning Toolbox (The MathWorks). Structural data matrices were mean-centered column-wise (default option for *plsregress*) prior to analysis. Predictor matrices were restricted to columns that had greater than two non-zero observations.

We fit PLSR models for each FC measure using the four different structural lesion measures as predictors. Leave-one-out (i.e. jackknife) optimization was used to identify the number of components (i.e. predictors) included for each model by adding components and measuring the change in prediction error with the inclusion of each additional component (Abdi, 2010). Components were added until the sum of squared prediction errors for the held-out cases increased with the addition of the new component, as increases in prediction error following the inclusion of additional components indicate overfitting to the training set (Abdi, 2010). This approach has been previously used for similar neuroimaging applications of PLSR (Kuceyeski et al., 2016b). The number of components for each model is shown in **Fig. 4B**.

PLSR models were then fit to the full dataset using the optimal number of components identified for each model (Kuceyeski et al., 2015, 2016b). Bootstrap resampling (1000 bootstraps) was used to estimate CIs for the model fits and beta weights using the bias-corrected and accelerated percentile method (Efron and Tibshirani, 1986) as implemented in the *bootci* function in MATLAB (**Fig. 4A, Fig. 5**). 95% CIs for model fits were adjusted to control the family-wise error rate for all 4 models fit to each FC measure, and therefore correspond to ∼99% confidence intervals. 99% CIs were also estimated for the beta weights from each model. The signs of model weights were flipped as necessary so that positive weights predicted more severe FC disruptions for all models. Beta weights were also rescaled for the plots in **Figure 5** and **Figures S4-S5** by multiplying all weights by a scalar value of 1000 (i.e. so that scientific notations would not overlap with the plot titles).

Comparisons of the different anatomical models of each FC outcome were performed using Akaike’s information criterion weights (AICw; **Fig. 4C**), as they incorporate information about both goodness-of-fit and model complexity (Kuceyeski et al., 2016b; Wagenmakers and Farrel, 2004). For each outcome variable, AICw were calculated as:

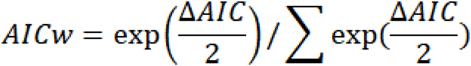

where Δ*AIC* corresponds to the difference between the AIC of each model and the minimum AIC across models for that outcome variable. The AIC weights for a given model from a set of candidate models can be interpreted as expressing the conditional probability that a given model is the best of all candidate models when considering both model performance and model complexity (Wagenmakers and Farrel, 2004). Thus, models with AIC weights closer to 1 are considered superior to models with AIC weights closer to 0. Linear correlations were computed among the unthresholded parcel SDC weights from all 12 models and among all 12 FC measures (**Fig. 5A**). Parcel SDC model weights were visualized using the SurfIce software package.

### Partial least squares correlations

We used partial least squares correlation (PLSC) of the full parcel-wise disconnection and FC datasets to identify the patterns of structural disconnection and FC that maximally covary across patients (**Fig. 6-8**). PLSC is a data-driven technique that is closely related to PLSR. PLSC seeks to define linear combinations of two data matrices (**X** and **Y**), referred to as latent variables (LVs), that maximally explain the covariance between the data matrices, and essentially involves performing a singular value decomposition (SVD) on the cross-block covariance matrix (Abdi, 2007; Krishnan et al., 2011; McIntosh and Lobaugh, 2004). PLSC has been successfully used to characterize covarying patterns of structural and functional connectivity in healthy individuals (Mišić et al., 2016) and has been successfully applied to other problems involving the relationships between structural and functional connectivity (Shen et al., 2015a, 2015b; Zimmermann et al., 2016).

Prior to performing the PLSC analysis, the upper triangle (excluding diagonal elements) of each patient’s *z*-transformed functional connectivity matrix was extracted and reshaped into a 1-dimensional vector. The resulting vectors were then stacked on top of each other to create a patient-by-edge FC matrix, and an analogous patient-by-edge matrix that was created using the parcel-wise disconnection matrices. Because the PLSC approach cannot accommodate missing values, FC between parcels that had been excluded from previous analyses (i.e. parcels with <60 vertices remaining) was set to 0 as in previously published multivariate analyses involving dense FC matrices from this sample (Siegel et al., 2016). However, analyses that were performed without removing lesioned parcels produced highly similar results (not shown), and control analyses that only included patients for whom no parcels were removed (*n*=51; see *Additional Analyses*) also produced results that were highly consistent with the main analyses (**Fig. S7**). The patient-by-edge parcel disconnection (**X** matrix) and functional connectivity (**Y** matrix) matrices were mean-centered column-wise and used to compute the covariance matrix **X’Y**. Singular value decomposition (SVD) was then applied using the MATLAB function *paq* to obtain the solution:

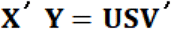

where

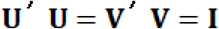

producing a set of *N*-1 orthogonal LVs that each consisted of singular vectors **U** and **V**, and a diagonal matrix **S** containing the singular values. The singular vectors contained weighted linear combinations of the original data matrices that maximally covaried together, and the singular values encoded the proportion of the covariance between the original data matrices that was accounted for by each LV. Score matrices were computed by multiplying the original data matrices by the corresponding singular vectors, analogously to PCA.

Permutation testing was used (1,000 permutations) to determine the significance of individual LVs (Abdi, 2007; Krishnan et al., 2011; McIntosh and Lobaugh, 2004), and bootstrap resampling (1,000 bootstraps) was used to compute bootstrap signal-to-noise ratios (BSRs) for the singular vector loadings associated with each LV by dividing the loadings by their bootstrapped standard error estimates (Abdi, 2007; Krishnan et al., 2011; McIntosh and Lobaugh, 2004). The (BSRs) quantify the stability of the loading estimates, and approximate z-scores (Efron and Tibshirani, 1986). Because the permutation and bootstrap procedures can produce LVs that do not match those obtained from PLSC of the original data, Procruste rotation was applied to the LVs obtained from the permutation/bootstrap analyses to ensure that they corresponded to those obtained from the original analyses (Krishnan et al., 2011; McIntosh and Lobaugh, 2004; Mišić et al., 2016). LVs obtained from the main PLSC analyses were considered significant if the permutation p-value was less than 0.05 after correcting for tests across all 10 LVs that accounted for at least 1% of the covariance, and loadings were considered stable if the corresponding absolute BSRs were greater than 2.5 (i.e. *p*∼0.01) (Krishnan et al., 2011; Mišić et al., 2016). Relevant results are shown in **Figure 6** and **Figures S6-S8**.

Linear correlation was used to assess the strength of the relationship between LV1 FC and disconnection scores. Bootstrap resampling (1,000 bootstraps) was used to compute a 95% confidence interval on the correlation (Krishnan et al., 2011). Patient scores on the first LV (LV1) were linearly correlated with mean interhemispheric within-system FC, ipsilesional DAN-DMN FC, and FC modularity measures, and with each of the behavioral measures (**Fig. 7**). Linear correlations were also computed between the unthresholded disconnection and FC loading vectors for LV1 to characterize the topographic similarity of the linked SDC and FC patterns (**Fig. 8**). This analysis only considered loadings that were non-zero in both vectors (i.e. only cortico-cortical edges that had disconnection loadings). FDR correction was used to correct for multiple testing for each set of correlations.

## Additional analyses

We performed additional analyses to (1) ensure that our main PLSR and PLSC results were not impacted by vascular factors as indexed by hemodynamic lags (Lv et al., 2013; Siegel et al., 2015), (2) ensure that our main PLSR results held when all models were fit with only a single component, (3) ensure that our main PLSR and PLSC results did not depend on the inclusion of patients with lesions in either hemisphere, (4) ensure that the topographical similarity analyses of the PLSC loadings were not driven by the exclusion of highly damaged parcels from the FC estimation, (5) ensure that the topographies of SDC PLSR model weights were not distorted by including only SDC information in the SDC PLSR models, and (6) ensure that the low-dimensionality of the PLSC results was not constrained by an intrinsic low-dimensionality of the structural and/or FC measures. These analyses are described in more detail below.

### Controlling for large ipsilesional hemodynamic lags in PLSR/PLSC analyses

Identification of patients with abnormal hemodynamic lags proceeded as follows. For each voxel, hemodynamic lags were with estimated with respect to the global GM signal using a window size of −8s to 8s (i.e. 4 TR), and the average difference in lag values between the lesioned and unlesioned hemispheres was computed for each patient as in previous work (Siegel et al., 2015). To ensure that our main results were not driven by patients with potentially abnormal hemodynamics, supplemental PLSR and PLSC analyses were performed that excluded patients with lag differences greater than 2 standard deviations from the control mean (i.e. > 0.32 s; 20/114 patients excluded). We note that this threshold (0.32 s) is conservative compared to thresholds used in prior work (Siegel et al., 2016; Siegel et al., 2018). Results from the PLSR and PLSC analyses that excluded high-lag patients were highly consistent with the main results and are shown in **Figure S3A** and **Figure S7A**, respectively.

### Controlling for differences in the number of PLSR components across models

The PLSR parcel SDC models presented in **Figure 4** often utilized more components than the damage models. While the number of components for each model was determined in a principled manner using jackknife cross-validation and the AIC weights incorporated information about model complexity, we wanted to ensure that similar results were obtained when all models utilized only a single component. We therefore fit all of the PLSR models using only a single component solution. The results of these analyses were highly consistent with the main analyses and are presented in **Figure S3B**.

### Separate PLSR/PLSC analyses of patients with left vs. right hemispheric lesions

The analyses presented in the main text utilized data from patients with lesion in either hemisphere. To determine whether our main results held when analyses were restricted to patients with lesions in a single hemisphere, we performed separate PLSR and PLSC analyses for patients with left vs. right hemispheric lesions. Results from the PLSR and PLSC analyses that were restricted to patients with lesions in a single hemisphere were highly consistent with the results from the main analyses and are shown in **Figure S3C-D** and **Figure S3F-G**, respectively.

### Controlling for removal of damaged parcels in PLSC analyses

As described in the description of the functional connectivity estimation procedures, lesioned vertices were not included in the estimation of functional connectivity, and parcels that had less than 60 vertices remaining after excluding lesioned vertices were removed for each patient by setting them to NaN. When computing functional connectivity summary measures, this allowed us to completely exclude highly damaged parcels. However, the PLSC analyses used the dense functional connectivity matrices and therefore could not accommodate NaN values. Therefore, functional connectivity for removed parcels was set to 0 as in previous multivariate analyses (Siegel et al., 2016). However, we were concerned that the removal of highly damaged parcels from the functional connectivity matrices might introduce systematic covariance between disconnections caused by the lesions that resulted in parcel removal and the zero-valued cells in the functional connectivity matrix, as this could bias the solution and lead to artificial topographic similarity between the structural and functional loadings. While parcel removals were relatively infrequent (**Fig. S7A**), we still wanted to control for this possibility. Therefore, we repeated the PLSC and topographic similarity analyses using only data from 51 patients for whom no parcels were sufficiently damaged to be removed from the analyses. These patients had essentially minimal to no cortical damage (**Fig. S7B**), and therefore the results could not be attributed to the effects of parcel damage on FC. The results obtained from these analyses were highly similar to those obtained from the main analyses and are presented in **Figure S7.**

### PLSR analyses with composite SDC and damage models

The results shown in **Figure S5** suggested that parcel damage PLSR models sometimes mis-localized WM damage effects to nearby GM parcels. This suggested that the parcel damage models were taking advantage of damage to GM parcels that was correlated with the WM damage effects identified by the voxel damage models. Because the parcel SDC models lacked explicit information about GM damage, we were concerned that the parcel SDC topographies might be susceptible to similar distortion. Therefore, we performed supplemental analyses to determine whether the parcel SDC weight topographies were affected by including information about parcel-wise GM damage. This analysis consisted of running the PLSR analyses with both the parcel SDC and parcel damage predictor measures as predictors in the same model, and then correlating the resulting (unthresholded) PLSR weight vectors with those obtained from the original analyses. This revealed that the weight topographies of the parcel SDC models were virtually unchanged by the inclusion of the parcel damage measures (across-model mean correlation of weight vectors = 0.98, SD = 0.04). However, this did have substantial effect on the parcel damage weight topographies (across-model mean correlation of weight vectors = 0.79, SD = 0.24), consistent with what would be expected under the scenario described above given that the FC measures were most strongly related to WM damage and SDCs.

### Dimensionality of lesion, SDC, and FC data

To ensure that the low-dimensionality of the PLSC results was not simply reflecting an intrinsically low-dimensionality of the structural or FC data, we performed principal component analyses (PCA) on the dense lesion, parcel SDC, and FC data from the patient sample (**Fig. S8**). Recall that in the PLSC analyses, the first 5 LVs explained 82% of the total covariance between the dense SDC and FC datasets (**Fig. 6A**). By comparison, the dense lesion data were relatively high-dimensional – 28 components are necessary to explain 80% of the total variance across all voxels in the lesion coverage zone. The dense parcel SDC data were lower-dimensional than the lesion data but were still relatively high-dimensional – 15 components were necessary to explain 80% of the total variance in parcel SDCs across all edges in the SDC coverage zone. Finally, the dense FC data were very high-dimensional – 72 components were necessary to explain 80% of the total variance in the FC data. These results indicate that there were far fewer salient covariance dimensions between the SDC and FC data than there were variance dimensions in the lesion, SDC, or FC data. This argues against an intrinsic low-dimensionality of the dense structural or FC data as a source of the observed low-dimensional covariance.

## Data and Software Availability

The full set of neuroimaging and behavioral data are available at https://cnda.wustl.edu/. Specific data and analysis scripts are available on request to the authors.

## Supplementary Material

**Figure S1.**
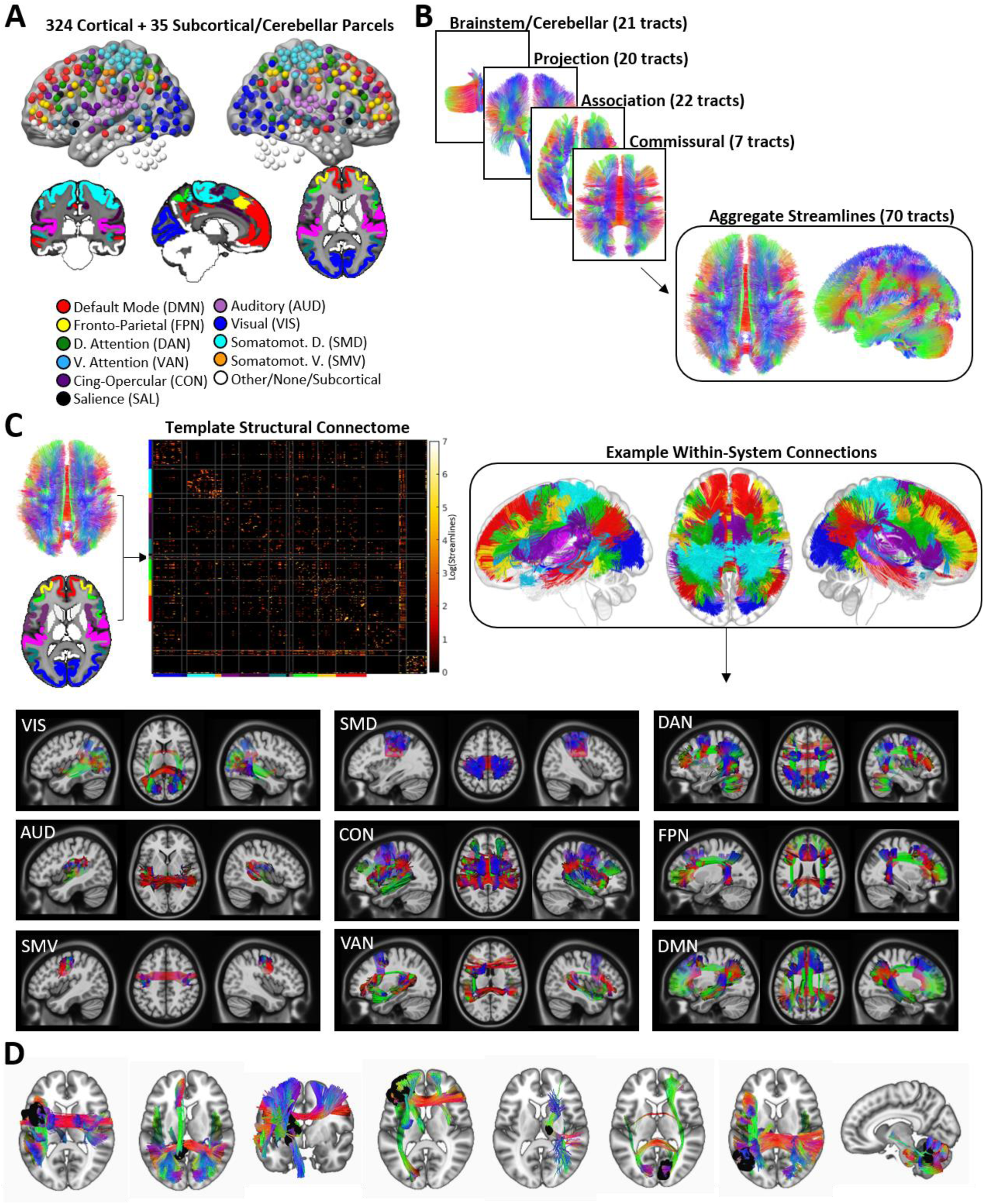
Related to Figure 1; Methods - Parcels and System Assignments, Structural Connectome Template. Parcels and templates. **A.** Spheres centered on the centroid co-ordinates of the 359 GM parcels are color-coded by system affiliations. Orthogonal slices below show volume-space parcels including subcortical and cerebellar parcels. **B.** Curated streamlines corresponding to 70 macroscale fiber pathways were combined into a single tractography atlas. **C.** The regional parcels and tractography atlas were used to construct a template structural connectome using endpoint-to-endpoint streamline extraction. The matrix shows the template structural connectome, and the colorscale corresponds to log(streamlines). The tractography image shows the within-system structural connections for the cortical systems shown in (A). Illustrations of structural connections for individual systems are shown below. **D.** Example disconnections for 8 patients with heterogeneous lesions. Lesions are shown in black.

**Supplementary Figure 2.**
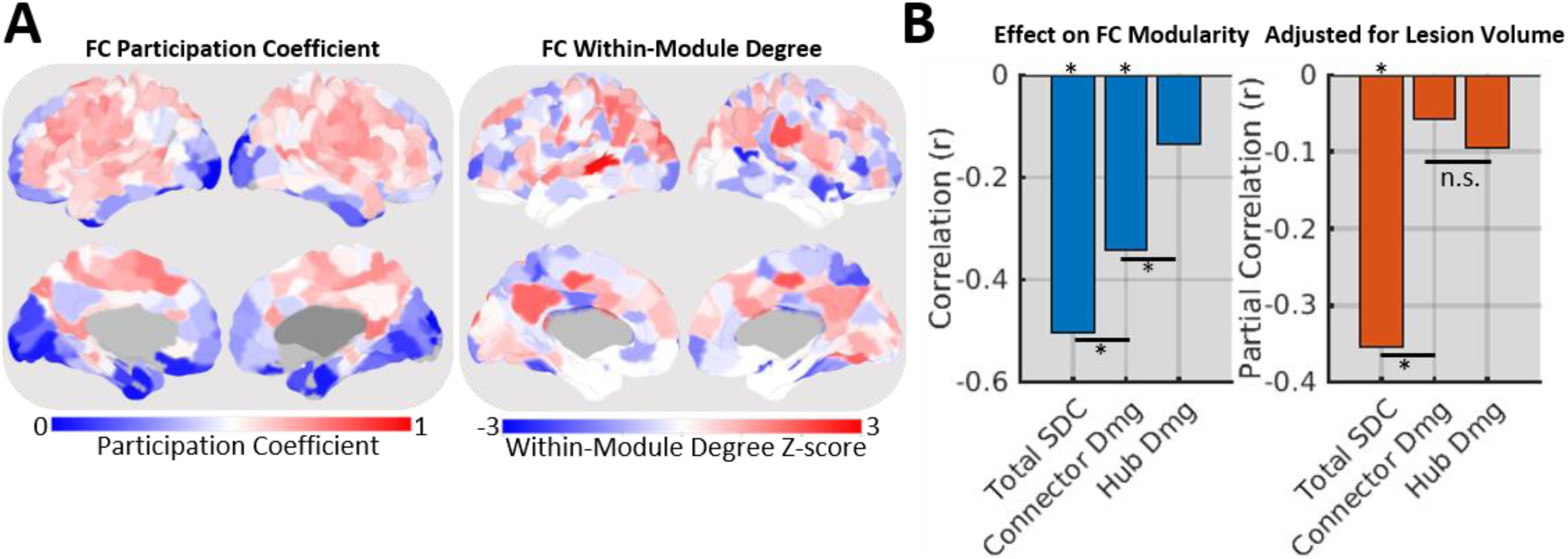
Related to Figure 3. Correlation-based analyses. **A.** FC participation coefficients (left) and FC within-module degree z-scores (right) for each parcel. Participation coefficients and within-module degree z-scores were estimated based on the mean FC matrix from the control group. **B.** The left plot shows the linear correlations between FC modularity and total SDC, FC connector damage, and FC hub damage. The right plot shows the partial correlations, adjusting for lesion volume. Steiger’s z-tests were used to compare effects between total SDC and FC connector damage (two-tailed), and between FC connector damage and FC hub damage (one-tailed – this effect was expected based on the results reported by Gratton et al., (2012)). *FDRp<0.05.

**Figure S3.**
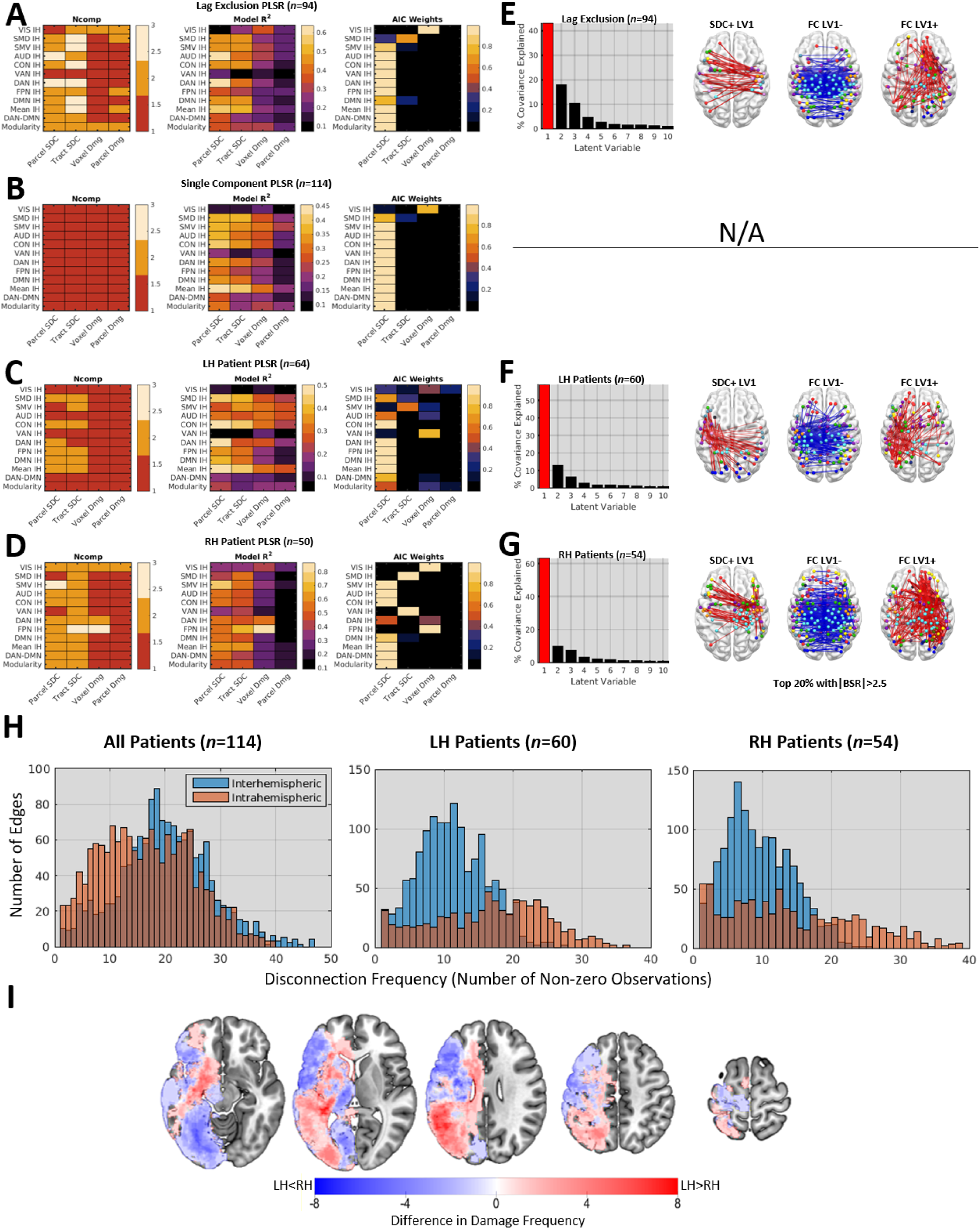
Related to Figure 4; Figure 6; Methods – Quantification and Statistical Analyses – Additional Analyses. **A-D.** Matrices show the number of components per PLSR model (left), PLSR model fits (middle), and PLSR AIC weights (right) from PLSR analyses where **(A)** analyses were restricted to patients with mean hemispheric lag differences less than 2 SD from the control mean (*n*=94), **(B)** analyses were run such that each model only included 1 component (*n*=114), **(C)** analyses were restricted to patients with left hemispheric lesions (*n*=60), and **(D)** analyses were restricted to patients with right hemispheric lesions (*n*=54). All analyses produced results that were highly consistent with the results obtained from the main analyses. Nearly identical results were also obtained even when lesioned vertices were included in the FC measures (not shown). **E-H.** The scree plots show the first 10 LVs obtained from PLSC analyses using the same criteria as the PLSR analyses shown in **(A-D)**. Note that there is no corresponding PLSC analysis for the 1-component PLSR analysis shown in **(B)**, and this is indicated by the “N/A” marking. The brain plots show the top 20% of significant PLSC loadings from each analysis. For the separate PLSC analyses of left and right hemispheric patients shown in **(F-G)**, the top weights for both groups featured a mixture of interhemispheric and intrahemispheric SDCs, along with FC patterns that closely resembled the FC pattern identified by the main analysis. **H.** The histograms show the number of edges (y-axes) with different SDC frequencies (i.e. number of non-zero SDCs; x-axes) for the analyses presented in the main text (left – All Patients), for the analyses of data from LH patients (middle – LH Only), and for the analyses of RH patients (right – RH Only). In each plot, separate histograms are shown for interhemispheric (IH – blue) and intrahemispheric (WH – orange) SDCs. The distributions of interhemispheric and intrahemispheric SDC frequencies were most comparable for the analyses presented in the main text, while intrahemispheric SDCs were more frequent than interhemispheric SDCs when analyses were restricted to patients with either left or right hemisphere lesions. This suggests that power to detect interhemispheric and intrahemispheric effects was most similar in the main analyses, while power was likely highest to detect intrahemispheric effects in the separate analyses. Further, effects that depended on lesion side (e.g. intrahemispheric SDC effects) were spatially independent in the main analyses, but not in the separate analyses. We consider it likely that the mixture of interhemispheric and intrahemispheric SDCs observed in the top loadings from the separate analyses reflects these factors. **I.** The map shows differences in voxel-wise lesion frequencies between LH and RH patient groups. The group-level lesion frequency map for RH patients was oriented to the left hemisphere and subtracted from the group-level lesion frequency map for LH patients to create the map. Blue voxels were damaged more frequently in patients with RH lesions, while red voxels were damaged more frequently in patients with LH lesions. Group differences in top SDC loading topographies shown in **(F-G)** largely reflect differences in lesion topographies between groups.

**Figure S4.**
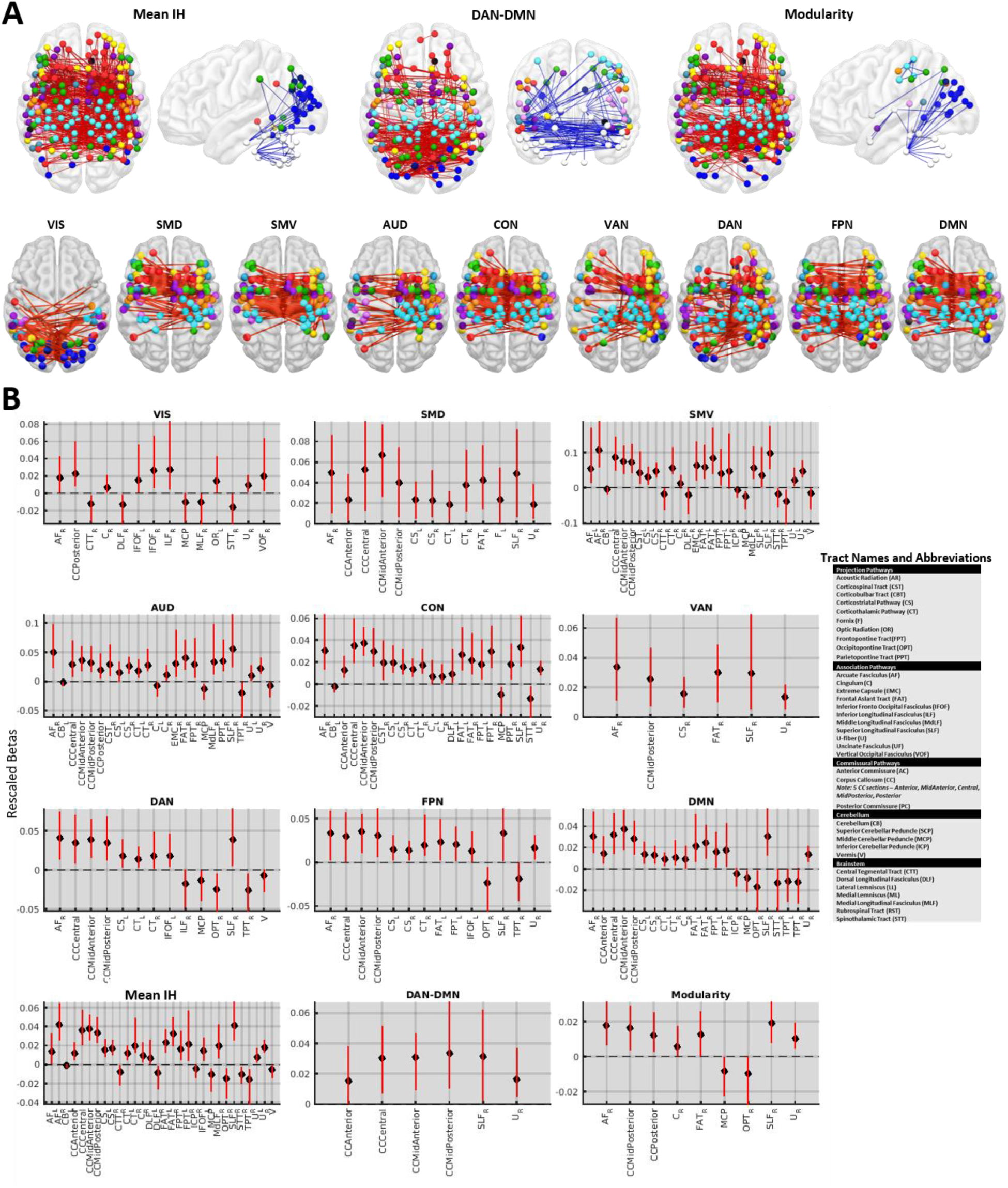
Related to Figure 5. PLSR SDC model weights. **A.** The top row shows the full PLSR weight topographies (i.e. all positive and negative weights with significant 99% CIs) for the three primary measures of interest (see **Figure 5**). The bottom row shows the top 20% of positive PLSR parcel SDC model weights with significant 99% CIs for the models of system-specific interhemispheric FC. Images include both interhemispheric and intrahemispheric disconnections. **B.** PLSR tract SDC model weights with significant 99% CIs are shown. Error bars correspond to 99% CIs as estimated via the bias-corrected and percentile-accelerated method. Tract names and abbreviations are provided in the legend. Weights are coded such that positive weights predict more severe disruptions of each FC measure, and negative weights predict less severe disruptions of each FC measure.

**Figure S5.**
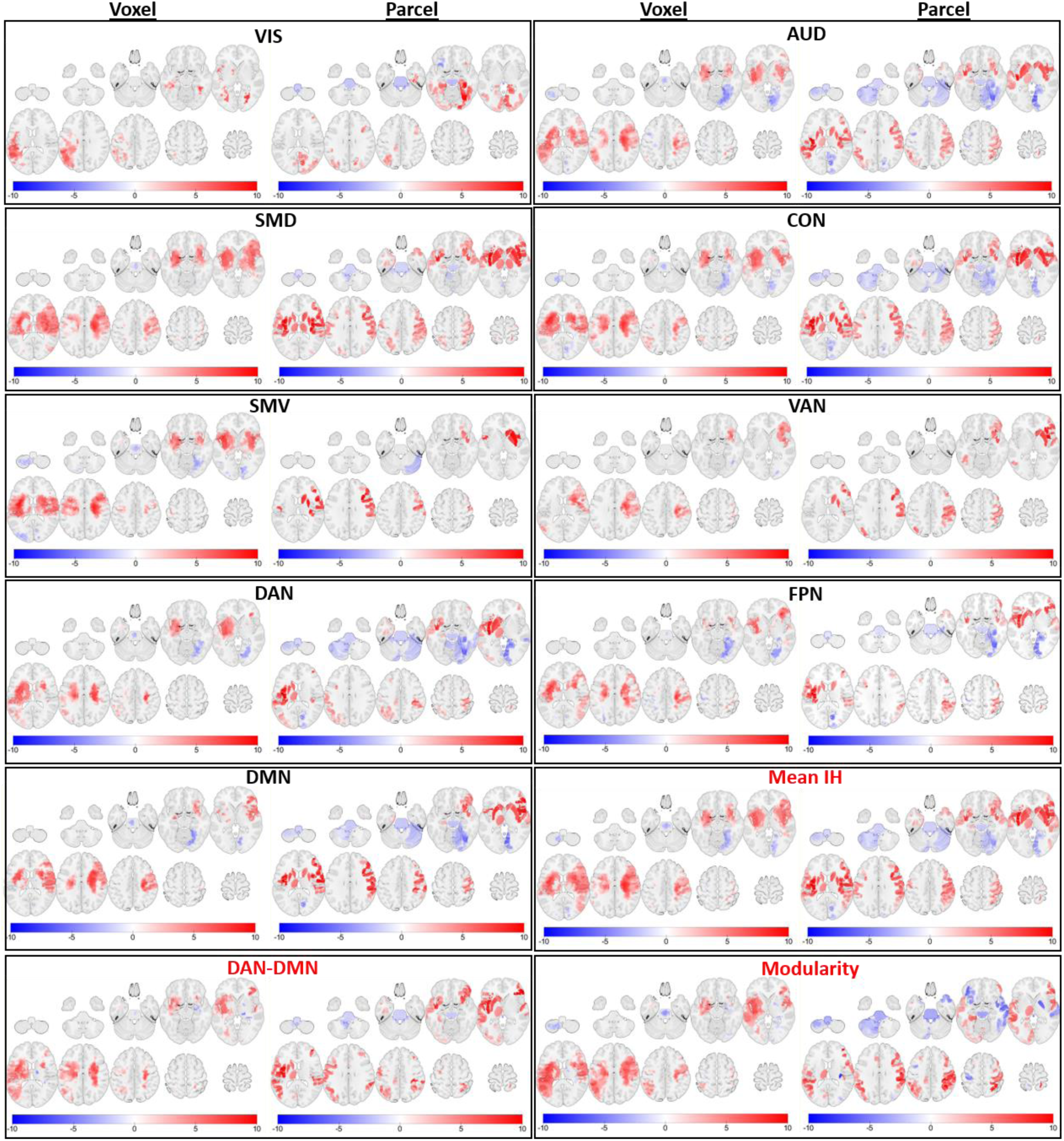
Related to Figure 5. Voxel and parcel damage PLSR model weights. PLSR voxel damage and parcel damage model weights with significant 99% CIs are shown for each model. Weights are coded such that positive weights predict more severe disruptions of each FC measure, and negative weights predict less severe disruptions of each FC measure. Weights for the models of the three primary measures are shown in the last three entries (names in red). Note that voxel damage maps emphasize white matter damage, and that white matter damage effects appear to be displaced into nearby cortical areas in the parcel damage maps.

**Figure S6.**
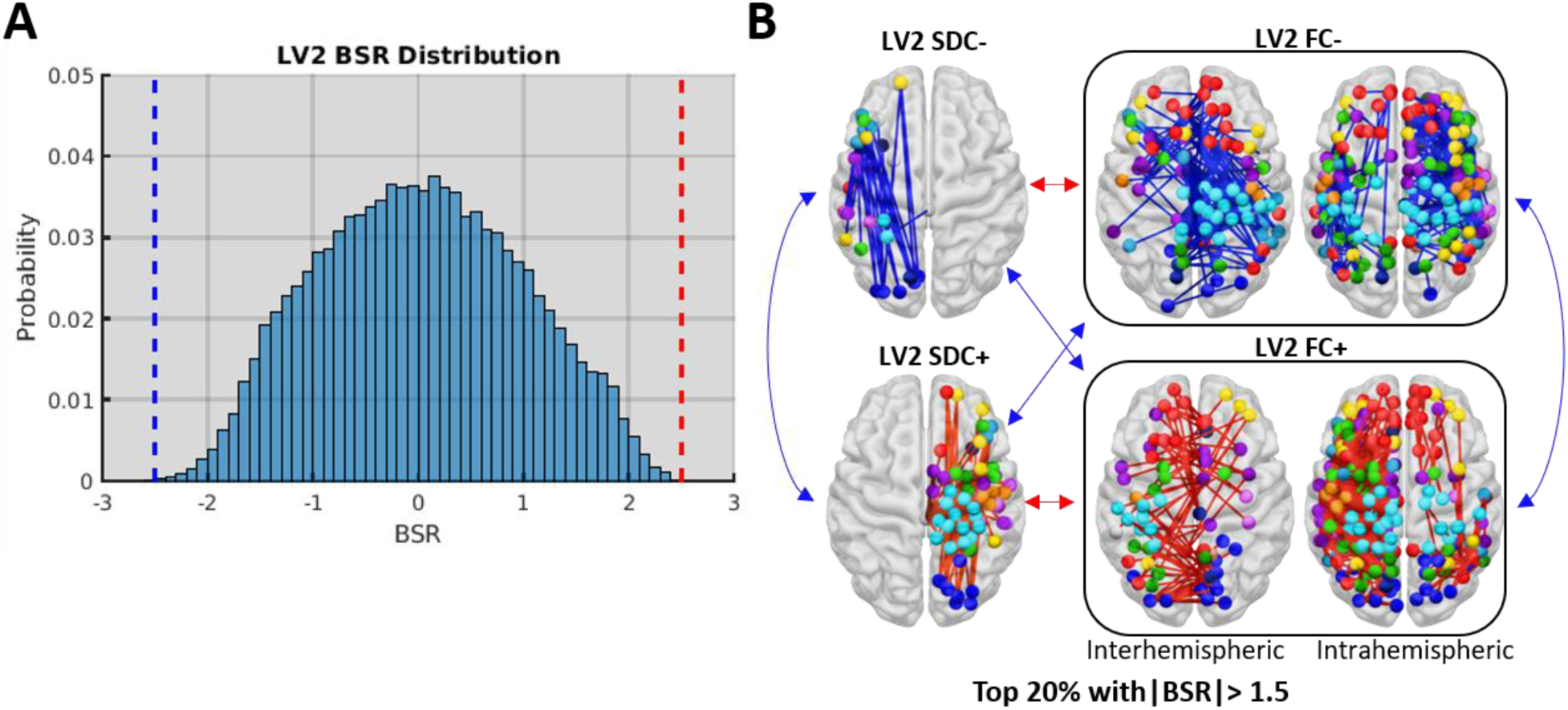
Related to Figure 6. PLSC LV2. A. The histogram shows the distribution of BSRs for LV2 (i.e. both SDC and FC BSRs). The dashed lines correspond to the significance thresholds (i.e. BSRs > 2.5, BSRs < −2.5) used in the main analyses. Loadings were not sufficiently stable to survive the significance threshold used for the main analyses. **B.** To illustrate the general patterns captured by LV2, the top 20% of positive and negative loadings with |BSRs| > 1.5 are shown. Arrows illustrate covariance relationships as in **Figure 6D**. SDC loadings on LV2 only included intrahemispheric edges, and loading signs differed for left vs. right hemispheric SDCs. FC loadings are shown separately for interhemispheric (left) and intrahemispheric (right) edges. FC loadings largely corresponded to intrahemispheric functional connections contralateral to SDCs with the same sign. Thus, LV2 appeared to primarily capture negative covariation between intrahemispheric SDCs and contralateral intrahemispheric FC. That is, less intrahemispheric SDCs within a given hemisphere were associated with stronger FC within that hemisphere, while more intrahemispheric SDCs within a given hemisphere were associated with weaker FC within that hemisphere. Interhemispheric functional connections largely featured midline regions and regions associated with somatomotor and default mode systems. Node system assignments are color coded as in **Figure S1**.

**Figure S7.**
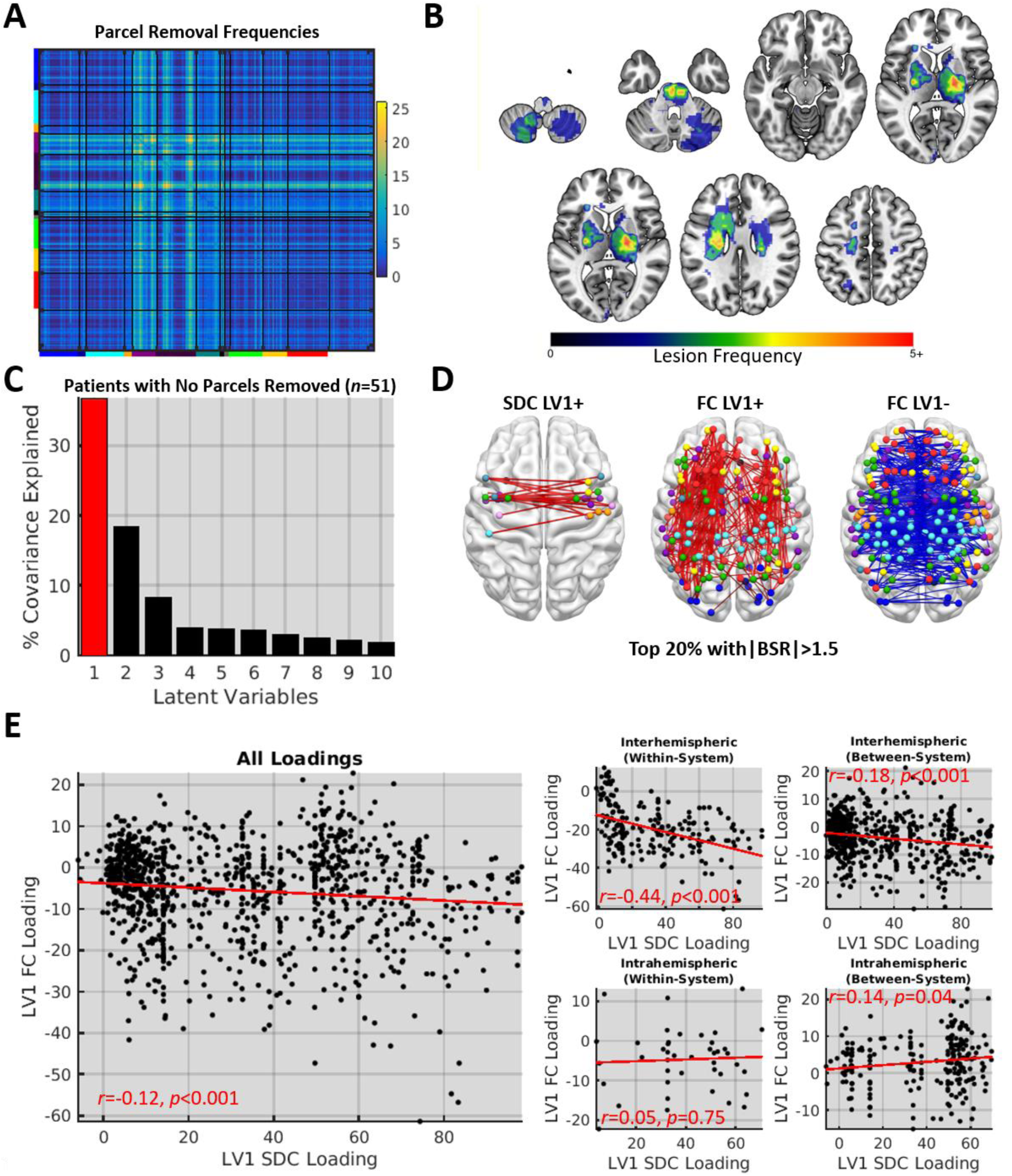
Related to Methods – Quantification and Statistical Analyses – Additional Analyses. Related to Figures 6 and 8. Results from PLSC analyses of patients with no parcels removed. **A.** The matrix shows the number of patients for whom each edge was removed. **B.** Lesion frequencies are shown for 51 patients who did not have any parcels that were sufficiently damaged to be removed. **C.** The scree plot shows the results of a PLSC analysis performed on the 51 patients with no parcels removed. **D.** The brain images show the top 20% of positive and negative SDC and FC loadings that survived a relaxed threshold of |BSR|>1.5 as very few loadings survived at the threshold used in the main analyses. **E.** The left scatterplot shows the relationship between all edges with non-zero LV1 SDC and FC loadings in the 51 patients with no parcels removed. The right scatterplots show the relationships between LV1 SDC and FC loadings for different connection types. While intrahemispheric within-system SDC and FC loadings were no longer significantly correlated, this was likely due to the small number of intrahemispheric within-system SDCs in this subsample. The significant correlations between interhemispheric/intrahemispheric between-system SDC and FC loadings may reflect the specific edges included in this sample. Despite these differences, these results argue that the topographic similarity between the SDC and FC loadings is not due to the removal of heavily damaged parcels from the FC matrices.

**Figure S8.**
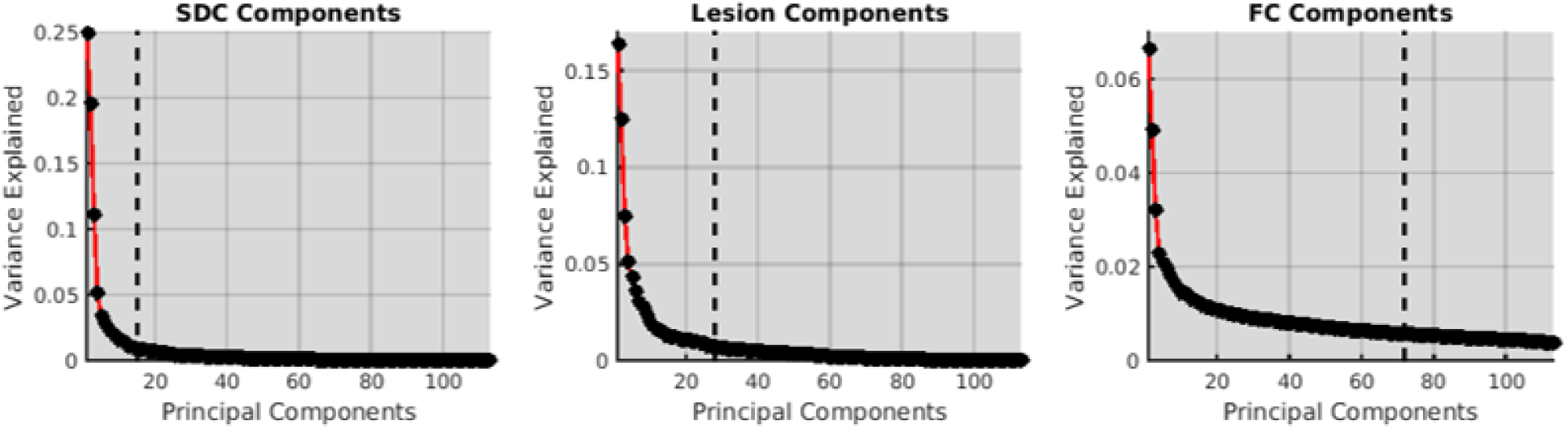
Related to Methods – Quantification and Statistical Analyses – Additional Analyses. Dimensionality of SDC, lesion, and FC data. Scree plots showing the proportion of variance explained by principal components of the parcel SDC (left), lesion (middle), and FC (right) data from the full patient sample (*n=114*). Dashed lines correspond to the number of components needed to explain 80% of the variance in each dataset. All datasets exhibited much higher dimensionality than the covariance between the SDC and FC datasets.

